# Characterization of TAG-63 and its role on axonal transport in *C. elegans*

**DOI:** 10.1101/723338

**Authors:** Prerana Bhan, Muniesh Muthaiyan Shanmugam, Ding Wang, Odvogmed Bayansan, Chih-Wei Chen, Oliver Ingvar Wagner

## Abstract

Model organisms are increasingly used to study and understand how neurofilament (NF)-based neurological diseases develop. However, whether a NF homolog exists in *C. elegans* remains unclear. We characterize TAG-63 as a NF-like protein with sequence homologies to human NEFH carrying various coiled coils as well as clustered phosphorylation sites. TAG-63 also exhibits features of NFL such as a molecular weight of around 70 kD, the lack of KSP repeats and the ability to form 10 nm filamentous structures in transmission electron micrographs. An anti-NEFH antibody detects a band at the predicted molecular weight of TAG-63 in Western blots of whole worm lysates and this band cannot be detected in *tag-63* knockout worms. A transcriptional *tag-63* reporter expresses in a broad range of neurons, and various anti-NFH antibodies stain worm neurons with an overlapping expression of axonal vesicle transporter UNC-104(KIF1A). Cultured neurons grow shorter axons when incubating with drugs known to disintegrate the NF network and rhodamine-labeled *in vitro* reconstituted TAG-63 filaments disintegrate upon drug exposure. Speeds of UNC-104 motors are diminished in *tag-63* mutant worms with visibly increased accumulations of motors along axons. UNC-104/TAG-63 and SNB-1/TAG-63 not only co-localize in neurons but also revealed positive BiFC (bimolecular fluorescence assay) signals.

## INTRODUCTION

The neuronal cytoskeleton consists of microtubules (MTs), actin filaments and neurofilaments (NFs). All of them are known to interact with each other providing the neuron a stable, polarized shape as well as important elastic features. Besides organelle positioning (e.g., nucleus, ER, and mitochondria), NFs are involved in intracellular signaling pathways and can be used as biomarkers ^1^. Heritable genetic mutations in NFs and the disintegration of the NF network are closely associated with various neurological diseases such as Giant Axonal Neuropathy, Charcot-Marie-Tooth (CMT) disease, Amyotrophic Lateral Sclerosis (ALS), and Parkinson’s disease (PD) ^2^. Besides the increasing knowledge on NF-based neuropathological diseases, little is known how such diseases develop on the molecular level. Model organisms are increasingly employed to study human diseases, including Zebrafish, *Drosophila* and *C. elegans*. Specifically, *C. elegans* is now widely used to study ALS, PD, Alzheimer’s disease (AD) and tauopathies ^3-5^. Noteworthy, it has been shown that NFL can be used as a biomarker for studying motor neuron degeneration along with tau protein ^6-8^. On the other hand, a NF-like protein has not been characterized in *C. elegans* yet, omitting this important model organism for studies on NF-based diseases. Thus, the major goal of this study was to identify and characterize a putative NF-like protein in *C. elegans*. Critically, increasing evidence suggests a role of axonal transport in a wide range of neurological diseases ^9,10^, and based on the fact that NFs are closely associated with MTs ^2^, we also intend to understand whether a putative *C. elegans* NF-like protein would affect axonal transport. While no specific neurofilament has been characterized in *C. elegans* yet, nematodes do express various intermediate filaments (IFs) which are categorized into eleven cytoplasmic IFs and one nuclear IF ^11^. These eleven cytoplasmic IFs are divided into an IFA/IFB-1 and an IFB-2/IFC/IFD/IFP-1-system. Obligate heteropolymers of IFAs and IFB-1 express in a diverse set of tissues, and they are essential for worm viability. IFs of the IFB-2/IFC/IFD/IFP-1-system mostly express in the worm’s intestine and they are non-essential for worm viability.

NFs in higher mammals are composed of three subunits, neurofilament-light polypeptides (NFL), neurofilament-medium polypeptides (NFM), and neurofilament-heavy polypeptides (NFH). Variations in the NF triplet protein have been identified in which the triplet incorporates either peripherin or α-internexin (depending on the cell type) ^12^. NF subunits possess highly conserved molecular domains including globular (N-terminal) heads, central coiled coil rod domains, and globular (C-terminal) tail domains. NFs extrude flexible tails (often termed “side arms”) to interact with other molecules such as MTs and organelles ^13,14^. NFH and NFM tail, but not NFL, polypeptides exhibit characteristic KSP (Lys-Ser-Pro) repeats often being targets for posttranslational phosphorylation modifications. Posttranslational modifications of NFs regulate their network integrity, organelle docking, and slow axonal transport ^15^. Although human NFL can homopolymerize to form filamentous structures ^16^, NFM and NFH can only heteropolymerize in the presence of NFL ^15^. NFs are insoluble (rope-like) filaments of 10 nm diameter that can be solubilized in the presence of urea to form intermediate oligomers ^17^. NFs undergo structural changes after urea treatment into globular units which can be renatured under prolonged dialysis against Tris-Buffer ^18^.

NFs often run in parallel to MTs, partially cross-linked via MAP2 ^19^, while NF side arms are able to cross-link NFs and to modulate interfilament spacing ^13^, thus regulating axonal caliber ^20^. NF sidearms are also critical to enhance interactions between NFs and mitochondria ^14^, as well as between NFs and dynein ^21^ or myosin Va ^22^. It has been shown recently in *C. elegans* that IFs hyperstabilize microtubules to prevent synapse rewiring consistent with the idea that IF levels are directly associated with axonal MT stability ^23^. Importantly, organelle transport is largely affected by IFs such as vimentin in non-neuronal cells ^24^, and in neurons the transport of NF subunits are tightly coordinated by kinesin-1 (anterograde direction) and dynein (retrograde direction) ^25^. It has been also observed that NFs alternate between two kinetic states, namely the mobile state and the stationary state during their transport along axons ^26^. NF subunits move in a stochastic, bidirectional and intermittent manner, supported by the “stop and go” transport model ^27^.

In *C. elegans*, kinesin-3 UNC-104(KIF1A) rapidly transports synaptic vesicles and their associated precursors in axons, such as SNB-1(Synaptobrevin-1), synaptotagmin, synaptophysin, and RAB-3 ^28^. Mice lacking KIF1A are embryonically lethal because synaptic vesicles are retained in cell bodies, leading to severe motor- and sensory neuron defects ^29^. Mutations in the *unc-104* gene result in paralyzed nematodes with vesicle retention phenotypes similar to mice ^30^. Overexpressing UNC-104::mRFP in worms with mutations in the *unc-104* gene fully rescues the paralyzed and highly uncoordinated *e1265* allelic phenotype ^31^.

The nervous system of *C. elegans* is simple (302 neurons), and specific types of neurons can be easily identified ^32^. Most motor- and mechanosensory neurons run along the worm’s body length while sensory neurons accumulate in the head with ciliated endings facing towards the “nose” of the worm. Head sensory neurons (e.g., amphid neurons) are often stabilized by supporting glial cells (sheath and socket cells) whereas their axons often inherit a short and ring-type appearance. Most somas of these head neurons accumulate in the nerve ring near the terminal bulb of the pharynx (a pumping foregut to aspirate nutriments) ^33^.

In this study, we attempt to identify and characterize a NF-like protein in *C. elegans* and to investigate its function in axonal transport. The outcome of such research might be critical to develop future worm disease models that may assist to understand the molecular nature of NF-based diseases.

## RESULTS

### Bioinformatic tools identify TAG-63 as a distant relative of human neurofilaments

Utilizing WormBase (wormbase.org) to identify NF homologs, we received a close hit for the gene *tag-63* (temporarily assigned gene-63) with BLASTP values of 5e-09 against human NEFH (or NFH, Neurofilament Heavy polypeptide) covering nearly all ten exons in TAG-63 (Suppl. Figure S1; note that to reproduce our results WormBase version number #WS230 must be used). Using KEGG database (kegg.jp), we received hits for NEFH orthologs from various animals (Figure 1B; Suppl. Figure S2), and further noted a cluster of NEFH orthologs (along with TAG-63) in a rooted phylogenetic tree emerging from a common ancestor (Suppl. Figure S2). Though we cannot detect KSP repeats in TAG-63, we identified nine (clustered) phosphorylation sites using the Scansite tool (scansite3.mit.edu) (Figure 1A, pink rectangles; Suppl. Figure S3A). Furthermore, the coiled coil prediction tool COILS/PCOILS (toolkit.tuebingen.mpg.de/pcoils) ^34^ identified three potential coiled coils in TAG-63 (Figure 1C). It needs to be mentioned that NIH BLASTP (blast.ncbi.nlm.nih.gov/Blast.cgi) does only deliver TAG-63 paralogs (Suppl. Figure S4C), however, employing NIH BLAST sequence alignment tool, we detected an average of 51% positives specifically located in the head and rod domains (Suppl. Figure S4A, and green boxes marked in S4B). We also attempt to investigate homologies between TAG-63 and various *C. elegans* IFs, and phylogenetic analysis (ClustalW2, ebi.ac.uk/tools/phylogeny) reveals IFP-1 as the closest protein (Suppl. Figure S5A), however, sequence alignment does only reveal poor similarities between TAG-63 and IFP-1 (Suppl. Figure S5B). In summary, bioinformatic tools revealed various sequence similarities between TAG-63 and NFH, while its low molecular weight (73 kD), the lack of KSP repeats with large phosphorylation-site cluster (highly phosphorylated NFs exhibit 40-51 phosphates per molecule) rather resembles NFL (also reflected in the phylogenetic tree by the distance between TAG-63 and other NFH homologs, Suppl. Figure S2).

**Figure 1:**
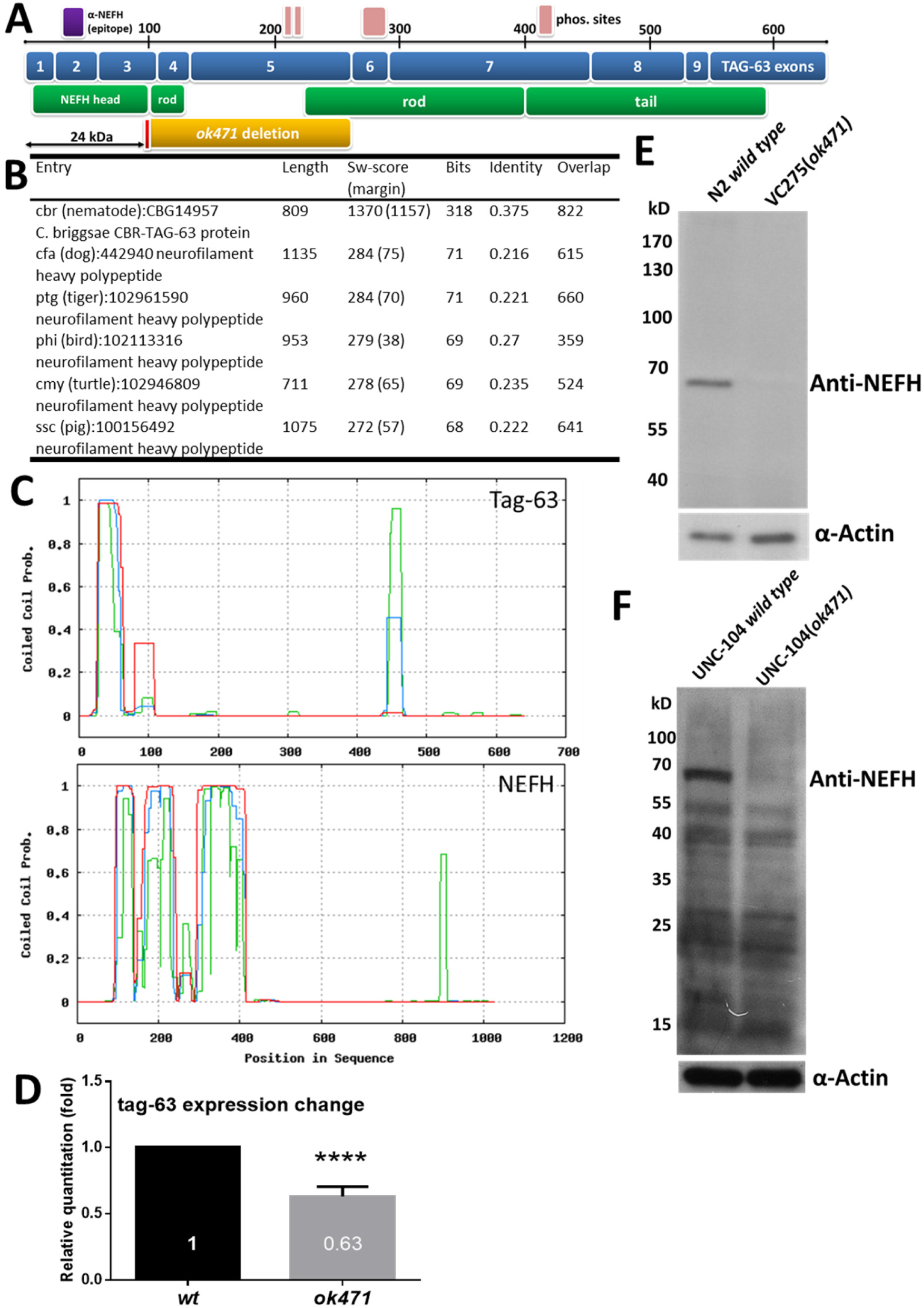
Identification and characterization of TAG-63 (Y47G6A.28). (A) The spliced *tag-63* gene covers ten exons (blue boxes) encoding for 641 amino acids with a calculated molecular weight of 73 kD. Matched similarities to human NEFH are indicated by green boxes and the matched epitope of the anti-NEFH antibody (#WH0004744M1, Sigma) is displayed by a purple box. The deletion region in *ok471* is indicated by a yellow box, and the identified insertion leading to a stop codon is indicated by a red box (likely resulting in a 24 kD truncated protein). Phosphorylation sites are represented by pink boxes. (B) KEGG database (kegg.jp) results with CELE_Y47G6A.28 query. (C) Coiled coil predictions of TAG-63 and NEFH (toolkit.tuebingen.mpg.de/pcoils). Green-window = 14 aa residues, blue-window = 21 aa residues and red-window =28 aa residues. (D) TAG-63 mRNA expression in wild type and *tag-63(ok471)* mutant quantified by Real-time PCR. Messenger RNA levels were normalized to *cdc-42* internal control. (E) Anti-NEFH antibody from mouse (#WH0004744M1, Sigma) detects a band around 70 kD which is not present in TAG-63 KO animals. (F) To understand whether a predicted truncated 24 kD TAG-63 product exist in mutants, a blot with low molecular weight marker is displayed. Here, we use lysates from worms expressing UNC-104::mRFP (used for studies as shown in Figure 6-9). Error bars: ± SEM. ****P<0.0001 (Student’s t-test).

### Characterization of TAG-63 in the *ok471* allelic background

Critically, a *tag-63* knockout strain VC275 (*ok471)* exist (generated by UV/TMP, ^35^), and the *ok471* allele has been annotated (wormbase.org) with a 1603 bp (264 aa) deletion eliminating exon 4 and 5 (Suppl. Figure S6A; amino acids of deletion area highlighted in blue in Suppl. Figure S6C). Our analysis also revealed a 51 bp insertion at position 194 in exon 3 (Suppl. Figure S6B green flanked region, and marked black in Suppl. Figure S6D with the first amino acid at position 194 marked in red) generating a nonsense mutation at position 203 (wildcard in Suppl. Figure S6B+D) that leads to a premature amber stop codon (likely generating a knockout). Real time PCR analysis revealed that *tag-63* levels were significantly reduced (up to 40%) in *ok471* mutants as opposed to wild types. This result also indicates that a quantity of truncated protein may be still produced in mutants (Figure 1D). A monoclonal anti-NEFH antibody (#WH0004744M1, Sigma) targets a region in exon 2 of TAG-63 (Figure 1A, purple rectangle, Suppl. Figure S3B). Using worm lysates from N2 wild type animals, we identified a band around 70 kD in Western blots (Figure 1E) close to the calculated 73 kD band. Importantly, in *tag-63* mutants, we cannot detect this band (Figure 1E) assuming that the strain VC275 (*ok471)* is a TAG-63 knockout (KO) strain. To understand whether or not a predicted deletion product of 24 kD (exon 1-3) is still produced in worms, we show another blot with a low molecular weight (MW) marker from a worm expressing kinesin-3 UNC-104 in *ok471* backgrounds (used for studies in Figures 6-9). Though we can identify a band around 24 kD in mutants (Figure 1F, UNC-104 (*ok471*)) this band is fairly weak and also appears in wild type animals (Figure 1F, UNC-104 wild type) indicating that either the 24 kD fragment is still weakly produced (though likely non-functional based on its truncated size) or this band reflects an unspecific band (frequently seen in whole worm lysates embodying more than 20,000 proteins).

**Figure 2:**
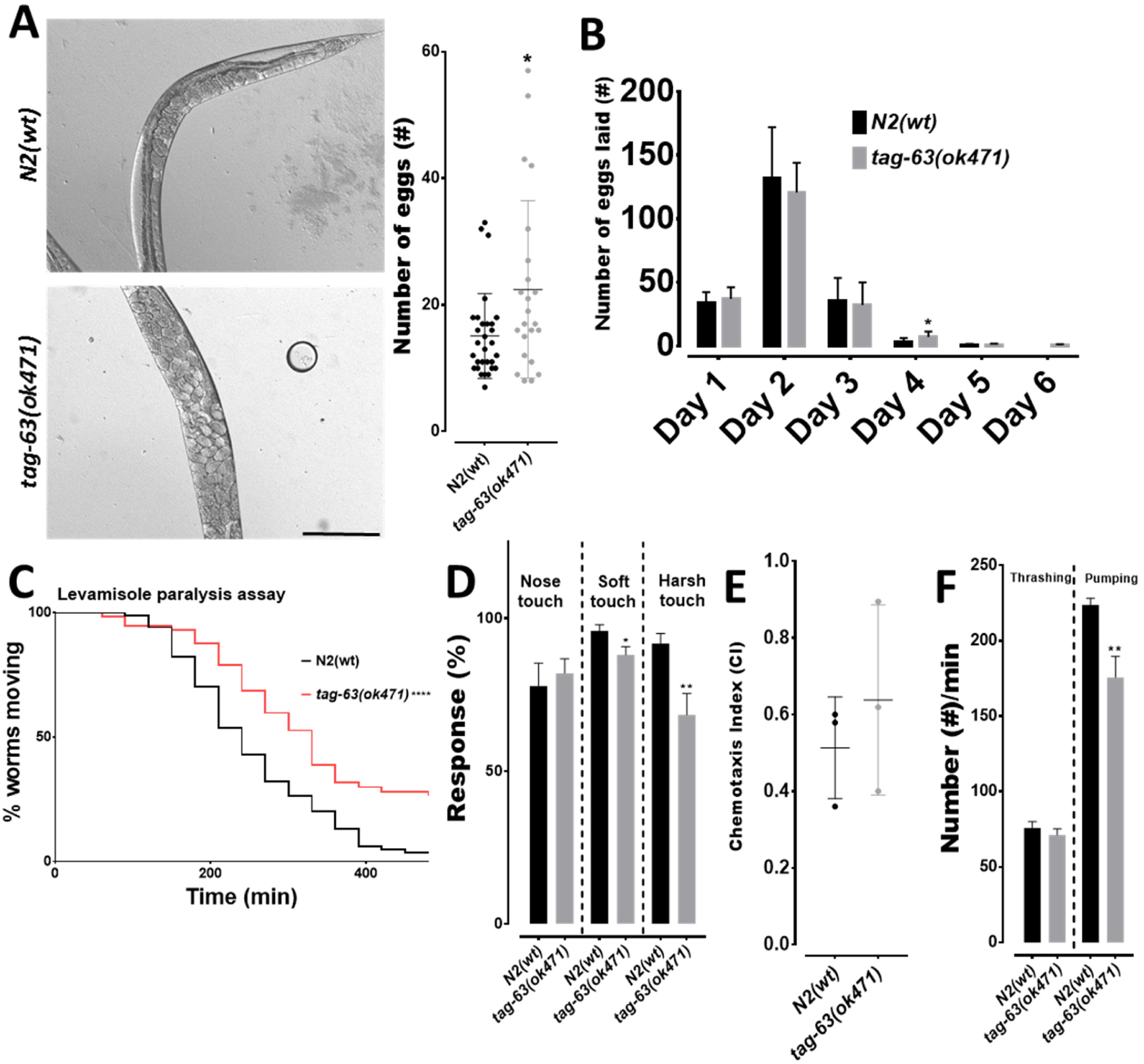
*Ok471* allele phenotyping. (A) Left: Bright field images of eggs retained inside the bodies of wild type and *tag-63(ok471)* mutant worms. Right: quantification, data points indicate number of eggs retained. (B) Eggs laid per day from the young adult stage (n = ∼10 worms). (C) Levamisole paralysis assay (n = 25/trial, total 3 trials). (D) Touch response analysis with nose touch (n = 12 worms), soft touch (n = 20-25 worms) and harsh touch (n = 20 worms). (E) Quadrant chemotaxis assay index, three independent trials with 19∼25 worms each. (F) Thrashing (n = 30 worms) and pharyngeal pumping frequency (n = 12 worms). Scale bar 100 µm. Error bars: ± SEM. *P<0.05, **P<0.01 and ****P<0.0001 (non-parametric Mann Whitney test (2A), Student’s t-test (2B, D-F) and log-rank test (2C)).

**Figure 3:**
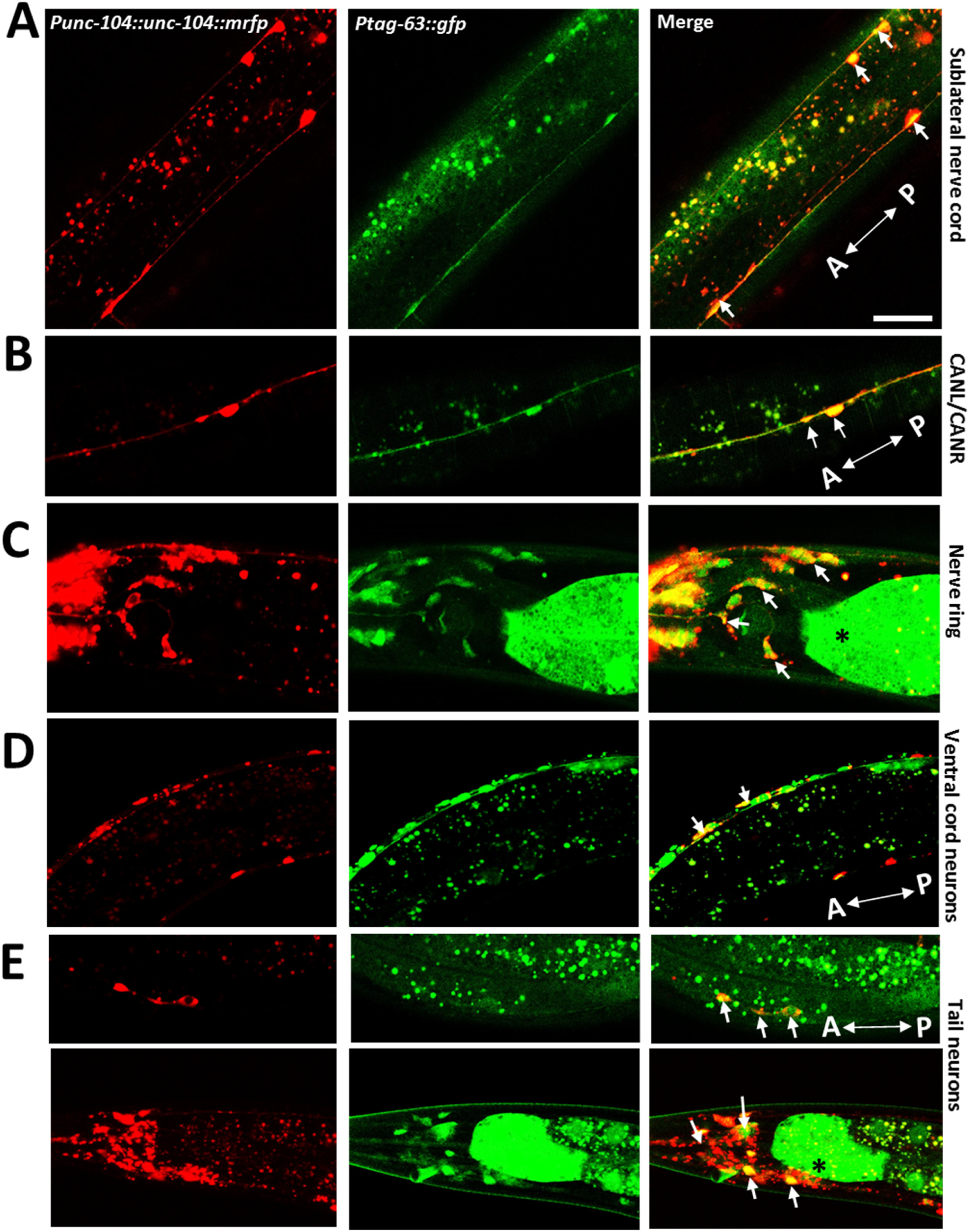
Characterization of *tag-63* expression in worms. Overview of a worm expressing *Ptag-63::gfp* reveals distinct expression in body- and head neurons (also see Suppl. Figure S7B-L). (A) *Ptag-63::gfp* expression in sublateral neurons that colocalizes with pan-neuronal *unc-104::mrfp* expression (for a simplified neuronal-network scheme in worms, see Suppl. Figure S7A). (B) Images revealing *tag-63* expression in CANL/CANR neurons, (C) *Ptag-63::gfp* expression in head neurons as well as in the nerve ring. (D-E) Expression pattern of *Ptag-63::gfp* in the ventral cord and tail neurons. White arrows indicate colocalization signals. Wildcards “*” in (C) and (E) denotes ectopic expression. A: anterior direction and P: posterior direction. Scale bar: 50 µm.

**Figure 4:**
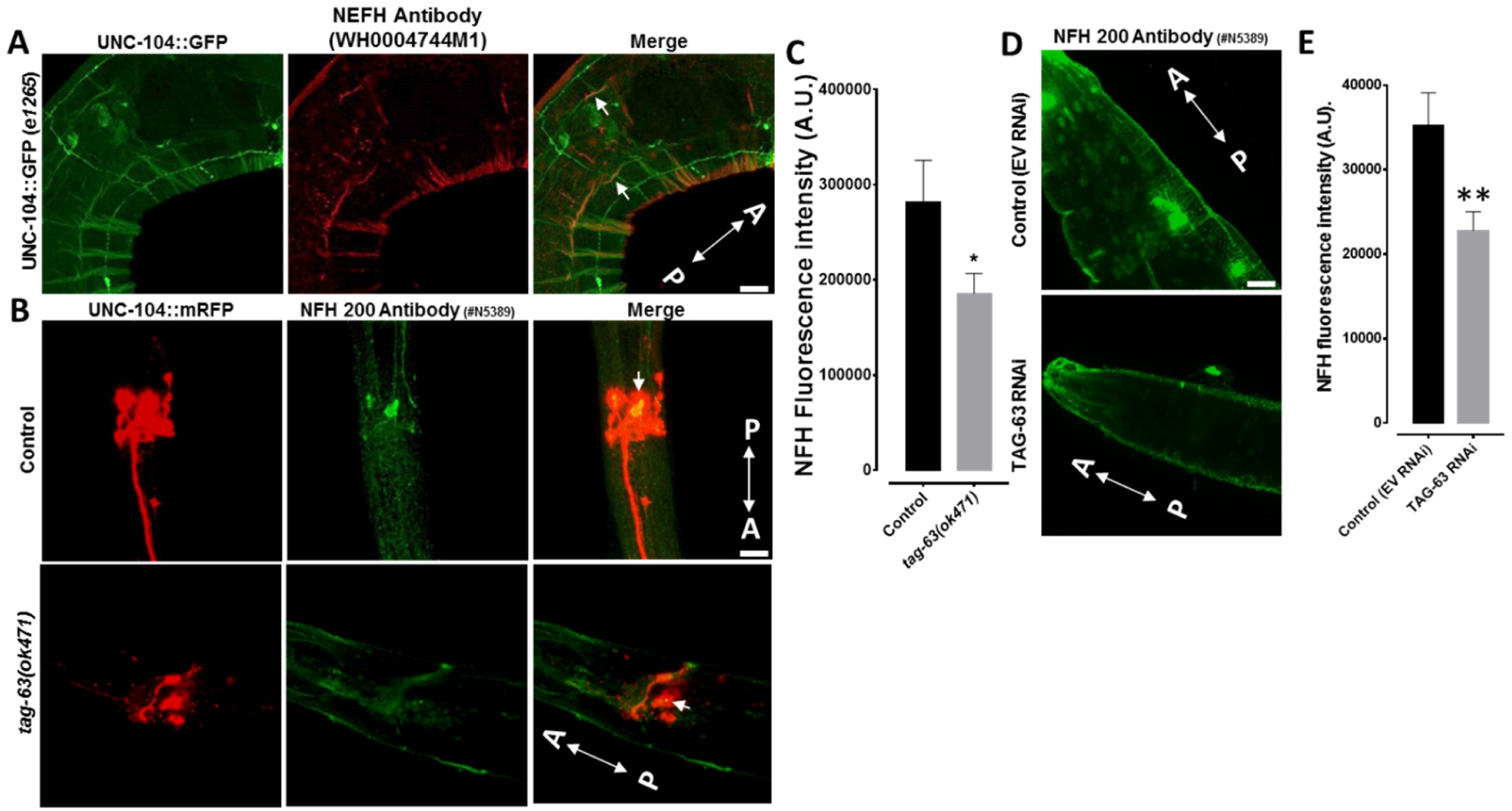
Whole mount immunostaining using various NFH antibodies. (A) Worm immunostained with NEFH antibody (WH0004744N1, Sigma). White arrow indicates staining in sublateral neurons overlapping with pan-neuronal *unc-104::gfp* expression. (B) Control and *tag-63(ok471)* mutant worms immunostained with NFH 200 antibody (#N5389, Sigma). White arrows indicate NFH staining in neurons overlapping with pan-neuronal *unc-104::mrfp* expression. Note that staining in *ok471* mutants is largely reduced compared to wild types (C). (D) Representative images of worms immunostained with NFH 200 antibody (#N5389, Sigma) and treated with either empty vector or TAG-63 RNAi. (E) Quantification of samples from (D) indicating a significant decrease in NFH staining after TAG-63 RNAi treatment. A: anterior direction, P: posterior direction. Scale bars: 10 µm. Error bars: ± SEM. *P<0.05 and **P<0.01 (Student’s t-test).

**Figure 5:**
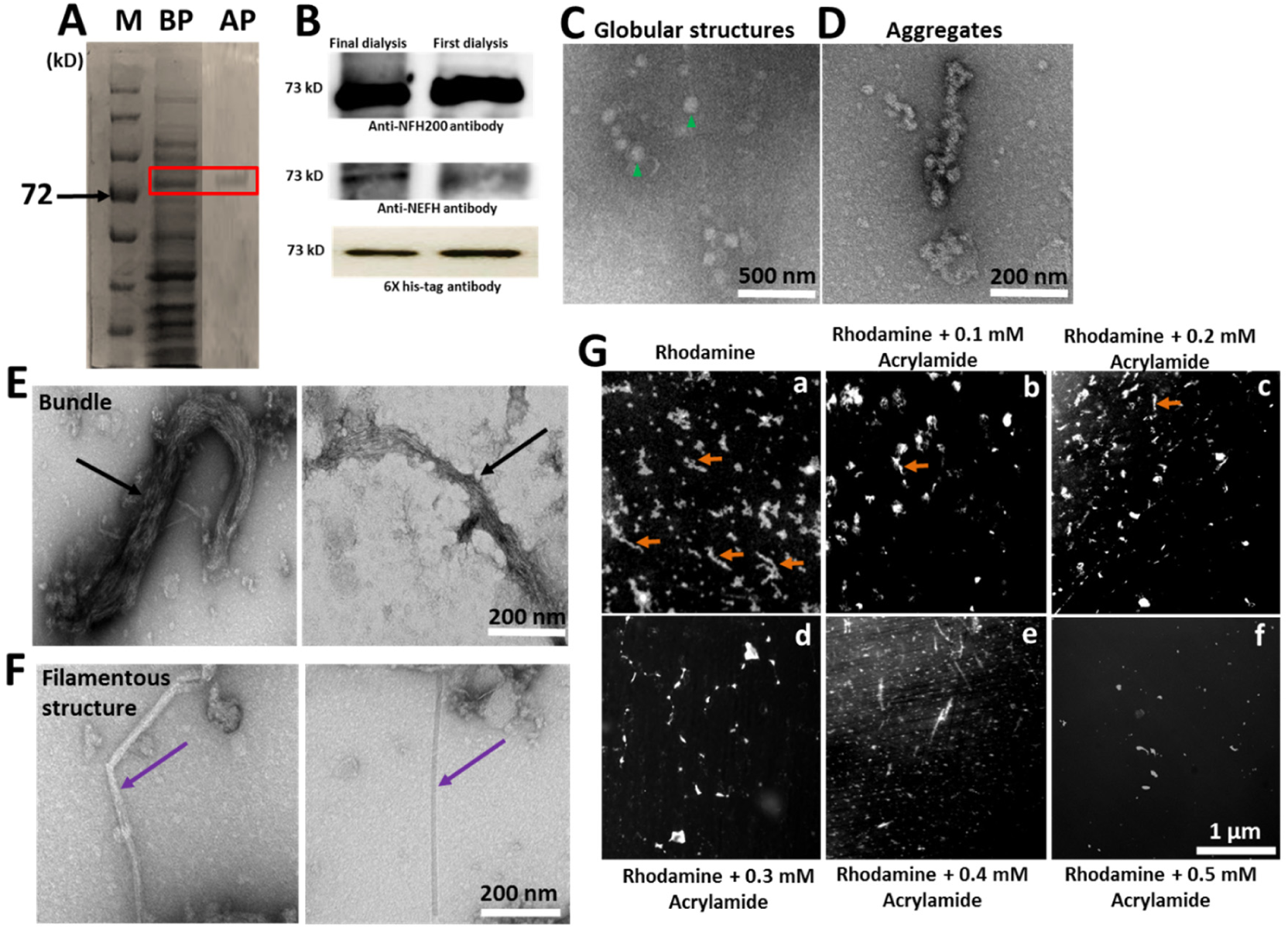
Electron micrographs and SDS-PAGE gels of purified TAG-63. (A) Commassie blue staining of proteins separated by SDS-PAGE before and after protein purification (marked by a red box). (B) Detection of purified TAG-63 protein at 73 kD by Western blotting using 6X His tag (GTX115045), anti-NEFH (produced in mouse, monoclonal, #WH0004744M1, Sigma), and anti-NFH 200 (produced in mouse, monoclonal, #N5389, Sigma) antibodies. (C-F) TEM images of negative stained purified TAG-63: After 8 M urea treatment TAG-63 appears as (C) globular entities (indicated by green arrow heads); while prolonged dialysis against 20 mM Tris-HCl buffer (pH 7.4) renaturation occurs into either, (D) aggregate-like structures, (E) loosely packed filamentous bundles (black arrow), or, (F) thin bundles with occasionally emanating single filaments of 10-12 nm in diameter (violet arrow). (G): (a) Fluorescence image of NHS-Rhodamine stained purified TAG-63. Orange arrows point to filamentous structures. (b-f) Rhodamine labeled TAG-63 treated with varying concentrations of acrylamide (0.1 mM - 0.5 mM). BP: before purification, AP: after column purification. Scale bars: (C) 500 nm, (D-F) 200 nm, and (G) 1 µm.

**Figure 6:**
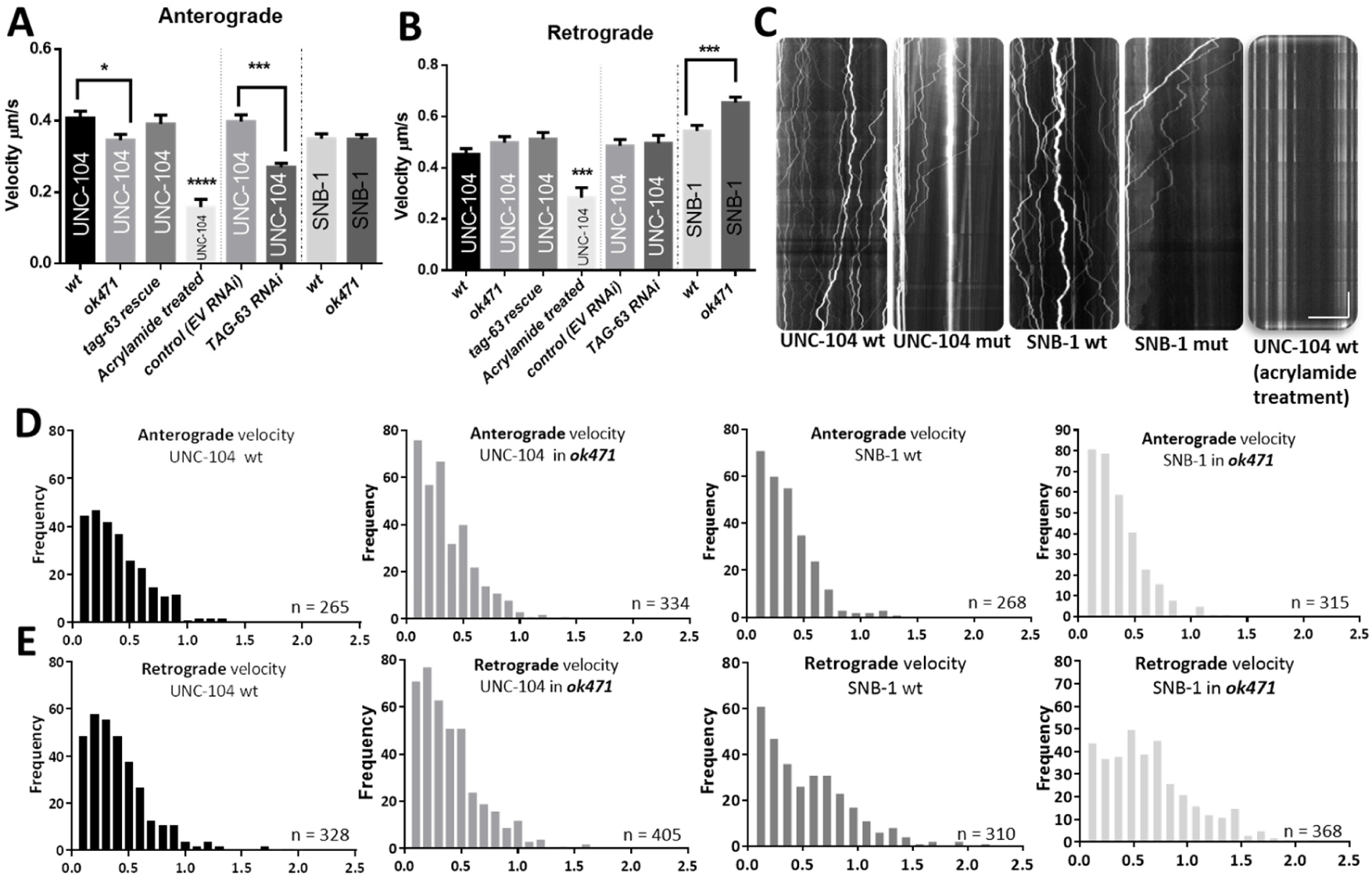
Effect of *tag-63* knockout on transport velocities of UNC-104 (motor) and SNB-1 (cargo). (A) In *tag-63(ok471)* mutants and TAG-63 RNAi knockdown animals, UNC-104 velocities are significantly reduced in anterograde directions, and this effect can be rescued by *tag-63::gfp* overexpression. Note that UNC-104 anterograde velocity is also markedly reduced after exposing worms to acrylamide. (B) Velocities of synaptic vesicles (marked by SNB-1) are significantly increased in retrograde directions when mutating TAG-63. Note the inhibitory effect of acrylamide on UNC-104’s retrograde velocity. (C) Representative kymographs used for motor (UNC-104) and cargo (SNB-1) motility analysis (wild type compared with *tag-63* mutant) as well as UNC-104 expressing worms treated with acrylamide. Anterograde direction is towards the right and retrograde direction is towards the left in the images. (D) Velocity distributions plotted in histograms for UNC-104 and SNB-1 in anterograde moving directions. (E) Velocity distributions for UNC-104 and SNB-1 in retrograde directions. Total analyzed events: UNC-104 in wild type = 593, UNC-104 in *ok471* = 739, UNC-104 *tag-63* rescue = 536, UNC-104 acrylamide treatment = 116, SNB-1 in wild type = 578, SNB-1 in *ok471* = 683, EV RNAi = 638, and TAG-63 RNAi = 526. “Mut” represents mutant *ok471*. Horizontal scale bar: 20 µm. Vertical scale bar: 20 sec. Error bars: ± SEM. *P<0.05, ***P<0.001 and ****P<0.0001 (Student’s t-test).

We then thoroughly investigated *ok471* phenotypes and detected egg retention phenotypes, reduced pharyngeal pumping rates, reduced responsiveness to soft and harsh touches as well as levamisole resistance (Figure 2A-D+F).

### Neuronal expression of TAG-63

To understand if *tag-63* is expressed in neuronal tissues, we co-injected a *tag-63* transcriptional GFP-reporter gene (*Ptag-63::gfp*) along with *unc-104::mrfp* into N2 wild type worms (Figure 3). Though the *tag-63* reporter occasionally reveals faint expression in gonads and intestines, we mostly detected strong expression in neurons (for a simplified neuronal-network scheme see Suppl. Figure S7A) such as body, head and tail neurons (Figure 3 and Suppl. Figure S7B-E). In detail, we identified significant expression in sublateral neurons (Figure 3A), CANL/CANR neurons (Figure 3B), nerve ring and amphid neurons (Figure 3C), ventral cord neurons (Figure 3D), tail neurons (Figure 3E) and mechanosensory neurons such as ALM (Suppl. Figure S7F-L). In many of these neurons TAG-63 co-localizes with (neuron-specific kinesin −3) UNC-104 and (synaptic precursor) SNB-1 proteins (Figure 9A-D).

**Figure 7:**
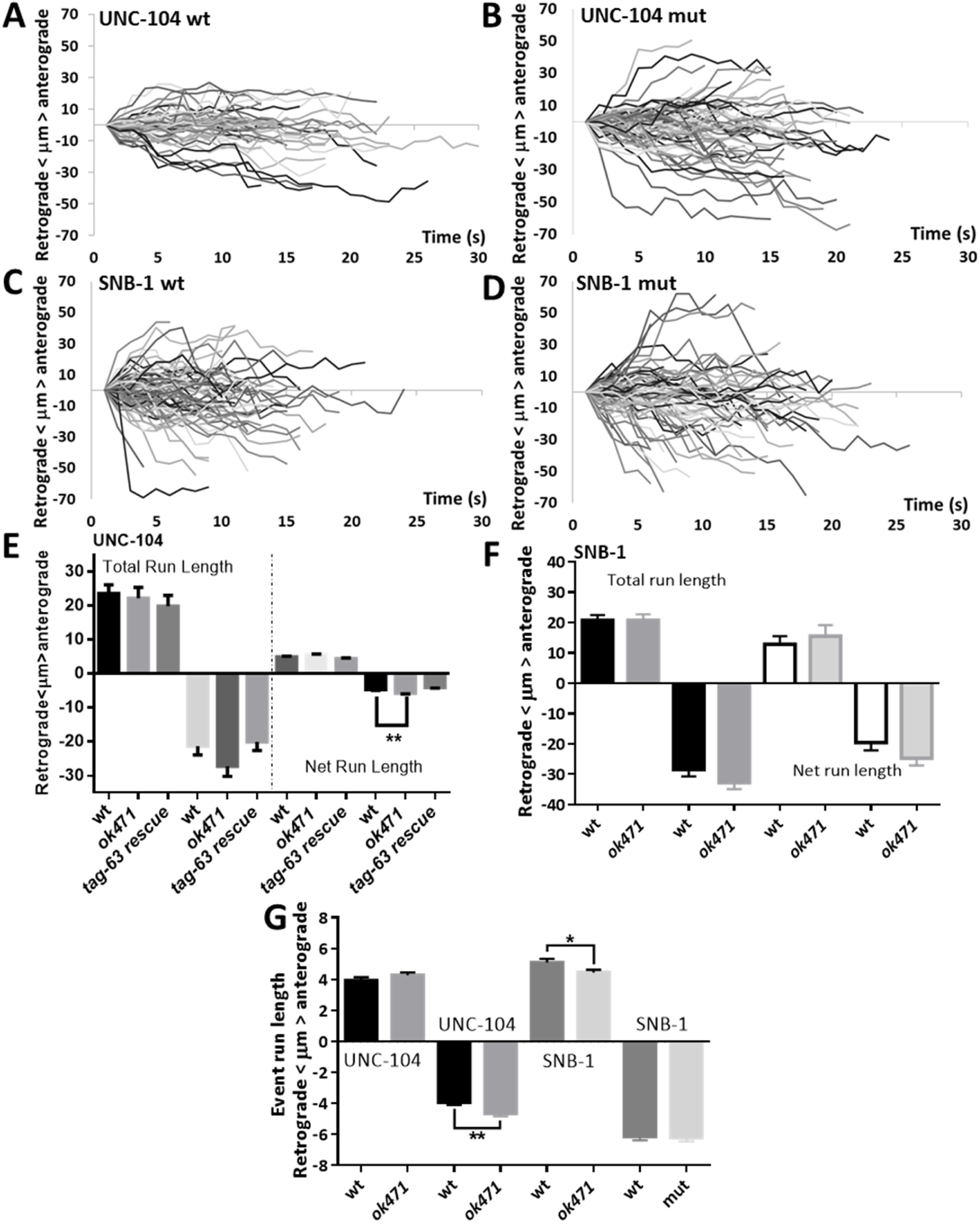
Effect of *tag-63* knockout on UNC-104 (motor) and SNB-1 (cargo) run lengths. (A-D) Traces from kymographs (i.e., Figure 6C) converted into bullet diagrams. (A) Bullet diagram of UNC-104 movement in wild type worms, and (B) of UNC-104 movement in *tag-63* mutant worms. (C) Bullet diagram of SNB-1 cargo in wild type worms, and (D) in *tag-63* mutant worms. (E) Average total run lengths and average net run lengths (for definition see Suppl. Figure S9D) in anterograde and retrograde directions of UNC-104 in wild type or *ok471* mutant’s. (F) Average total run lengths of SNB-1 in wild type (solid black bars) or in *ok471* mutants (solid gray bars). Average net run lengths are represented by open white bars for wild type and open gray bars for *ok471* mutants. (G) Single event run lengths in the anterograde and retrograde directions measured for UNC-104 and SNB-1 motility, respectively. Error bars: ± SEM. *P<0.05 and **P<0.01 (Student’s t-test).

**Figure 8:**
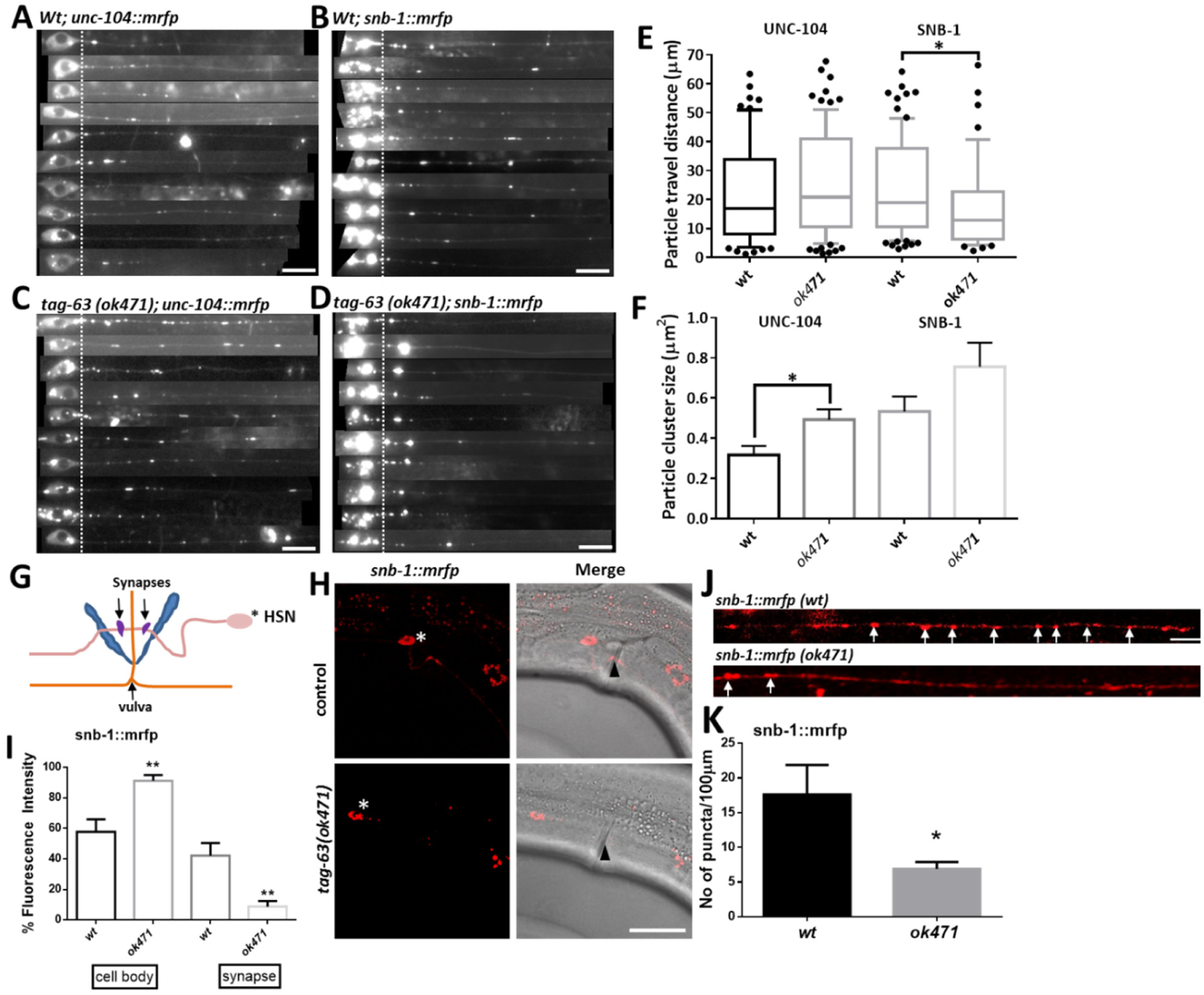
Particle distribution of UNC-104 and SNB-1 as well as targeting of SNB-1 to the synapse (*wt* compared to *tag-63*). (A+B) UNC-104 and SNB-1 particle distribution in ALM neurons in wild type worms. (C+D) UNC-104 and SNB-1 particle distribution in ALM neurons in *tag-63* mutant worms. (E) Distances of motors and cargos traveled from the proximal axon hillock (marked as dotted lines in A-D) to their final distal destinations (at the time of imaging). (F) Sizes of clustered UNC-104 and SNB-1 particles in wild type and *ok471* mutant worms, respectively. (G) Diagram representing wild type morphology and synaptic patterning in HSN neuron. HSN neuron that form synapses along vulva muscles are indicated in purple (wildcard denote soma). (H) *snb-1::mrfp* expression in soma and synapses of HSN neurons (wild types and mutants). Arrowheads point to vulva and wildcards indicate the location of somas. (I) Quantification of *snb-1::mrfp* fluorescence intensity in the somas and synapses of HSN neurons as shown in (H). (J) Localization of *snb-1::mrfp* in sublateral nerve cord. (K) Quantification of SNB-1 puncta number per 100 µm from (J). Scale bars: 10 µm. Error bars: ± SEM. *P<0.05 and **P<0.01 (Student’s t-test).

**Figure 9:**
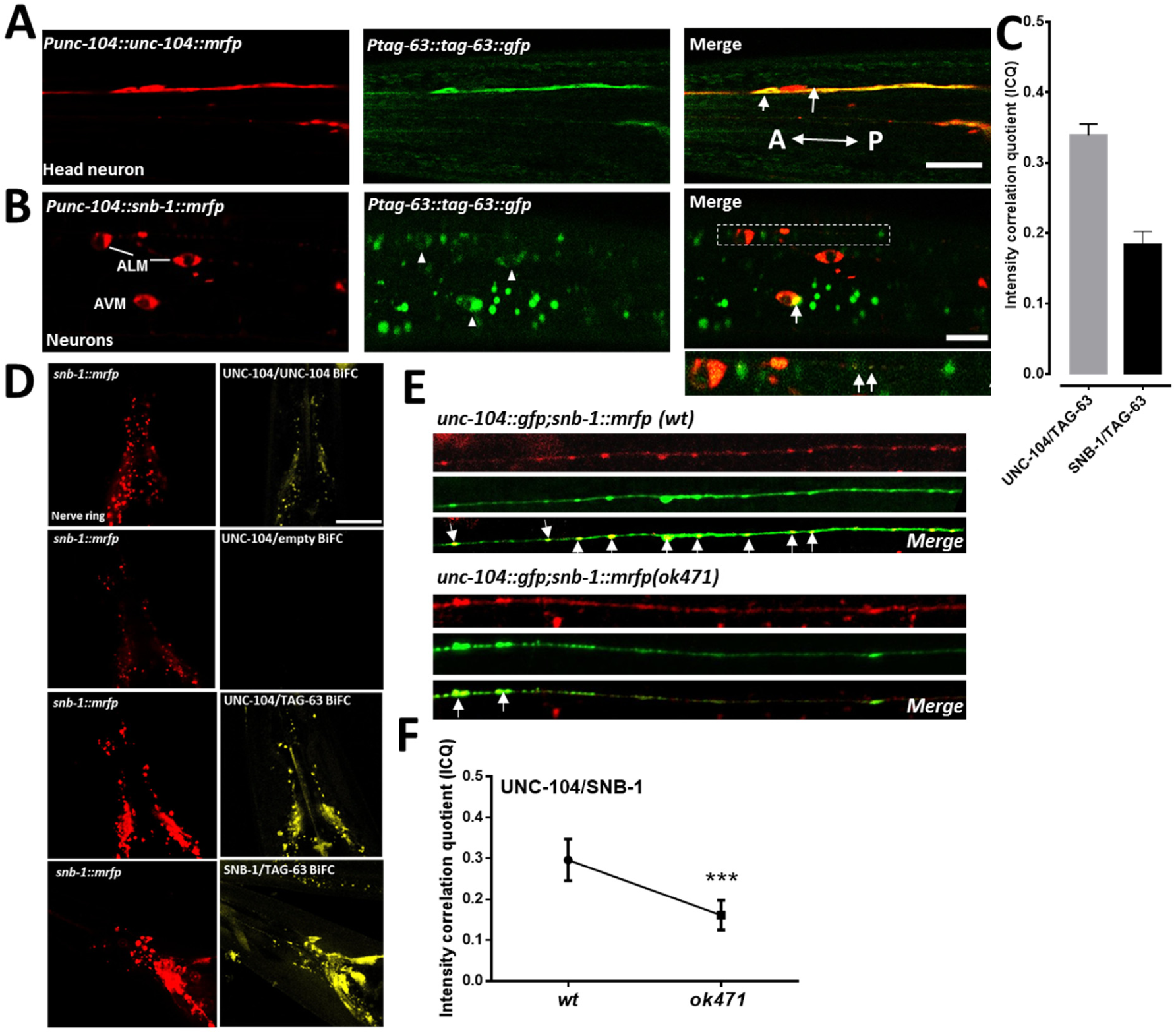
Colocalization of UNC-104 and SNB-1 with TAG-63. (A) Representative images of *unc-104::mrfp* colocalizing with *tag-63::gfp* in head neurons. (B) Images of *snb-1::mrfp* colocalization with *tag-63::gfp* in various neurons. A: anterior direction, P: posterior direction. White arrows (A+B) indicate colocalization signals and arrowhead (B) indicates somas. Dashed white rectangle indicates the magnified inset below. (C) Quantification of colocalization between both UNC-104/TAG-63 and SNB-1/TAG-63 using ICQ method. (D) BiFC signals from UNC-104/UNC-104 interaction (positive control). No BiFC signals were observed for UNC-104/empty vector interactions (negative control). BiFC signals from UNC-104/TAG-63 and SNB-1/TAG-63 interactions in nerve ring. (E) Colocalization of UNC-104 and SNB-1 in wild type and *tag-63(ok471)* mutant worms in sublateral nerve cord. (F) Quantification of UNC-104/SNB-1 colocalization (from (E)) using ICQ method. Error bars: ± SEM. Scale bars: 10 µm. ***P<0.001 (Student’s t-test).

Whole mount immunostaining employing either anti-NEFH ((#WH0004744M1, Sigma) and anti-NFH200 (#N5389, Sigma) antibodies all display specific staining in various neurons, and we confirmed colocalization between these fluorescently-tagged antibodies and pan-neuronal UNC-104 expression (Figure 4A-E). We also observed colocalization of NFH200 antibody staining (seen in cyan) with TAG-63::GFP and UNC-104::mRFP (Figure S8C+D; line scan represents overlapping fluorescence signals). Though the NFH200 antibody staining was significantly reduced in *tag-63(ok471)* mutant worms, we occasionally detected faint staining in some neurons, likely being the result of (non-degraded) truncated TAG-63 protein (compare Figure 4B+C, to 1D). Nevertheless, antibody staining after *tag-63* knockdown was significantly reduced (Figure 4D+E). Note that we have previously shown the efficiency of RNAi in neuronal tissues without the need of cross-breeding with RNAi sensitive strains ^36^, and also for this study we provide data reflecting the efficiency of *tag-63* knockdown in neurons (Figure S8A+B). Though we have no information regarding the epitope sequence of the NFH200 antibody, we obtained the immunogen sequence for the NEFH antibody, and our analysis revealed that this antibody may cross-react with TAG-63, but likely not with IFP-1, IFB-1 or IFA-4 (Suppl. Figure S3B).

### Ultrastructural studies reveal the ability of TAG-63 to form filamentous structures

It has been reported that acrylamide and TCP (tricresyl phosphate) affects structure and function of NFs ^37-39^ and we used cultured neurons from worm embryos (expressing pan-neuronal reporter UNC-104::GFP) to understand whether these chemicals would affect development and morphology of these neurons (note that we used cultured neurons to warrant proper uptake of these drugs). Indeed, incubating these primary neurons with either acrylamide or TCP resulted in neurons with shorter axons (Suppl. Figure S9A+B). Interestingly, these chemicals also affected the distribution of UNC-104 motors resulting in enhanced clustering along axons. We assume that a disintegrated axonal cytoskeleton (to which NFs largely contribute) ^40^ results in both shorter axons as well as motor accumulations. To understand whether TAG-63 indeed would form filamentous structures, and whether the formation of filaments would be negatively affected by acrylamide, we recombinantly expressed and purified TAG-63 protein. Figure 5A+B shows the band of the purified protein at calculated 73 kD identified by anti-His tag, anti-NEFH and anti-NFH200 antibodies (note that the protein size is slightly higher compared to that revealed by Western blot analysis in Figure 1E+F likely based on the additional linkers, T7 tag and His tag, in the expression vector). TAG-63 protein appeared as globular structures after urea solubilization of sizes ranging from 120 nm to 180 nm, respectively (Figure 5C), and these structures largely resemble those published for reconstituted NFL in TEM ^18^. As expected and reported earlier for NFs ^18^, these globular structures are able to convert into either unusually thick aggregates (Figure 5D) or fibrous bundles containing single filaments of 10 nm sizes (Figure 5E+F) after dialyzing against 20 mM Tris-buffer; and all of these structures (Figure 5D-F) largely resemble those published previously for *in vitro* reconstituted NFL ^18,41^. Furthermore, we stained purified TAG-63 protein with NHS-Rhodamine and identified filamentous structures using fluorescence microscopy (Figure 5G (a)) that resemble those shown previously for rhodamine-stained NFs ^14^. Moreover, we examined whether acrylamide (known to disrupt NFs, ^42^) would reduce the amount of filamentous TAG-63, and indeed we observed dose-dependent collapsing of TAG-63 filaments (to globular structures) with increasing acrylamide concentrations (Figure 5G(b-f)).

### Effect of *tag-63* on UNC-104 (motor) and SNB-1 (cargo) transport efficiencies

As the disintegration of both vimentin and NFs are known to affect MT-based transport ^43-45^, we analyzed the motility of kine-sin-3 UNC-104 and its cargo SNB-1 (synaptobrevin-1) in ALM neurons of wild type, TAG-63 knockout as well as TAG-63 knockdown (RNAi) worms. Figure 6 shows anterograde and retrograde velocity profiles of motor UNC-104 as well as of its cargo SNB-1 ^31,36,46^, both tagged with mRFP (revealed from separate strains and experiments). Anterograde movements of UNC-104 are reduced in *tag-63*(*ok471*) animals (Figure 6A+D) pointing towards a regulatory role of TAG-63 on UNC-104’s motor activity. Importantly, this effect can be rescued by overexpressing TAG-63 in *tag-63*(*ok471*) mutant worms (Figure 6A). Also knockdown of TAG-63 in UNC-104::mRFP*(e1265)* worms revealed significant reduction in anterograde velocities (Figure 6A). Interestingly, *tag-63* significantly affected the retrograde movements of SNB-1 and this effect is likely contributed by increasing dynein activity (Figure 6B+E). Though understanding the mechanistic basis of these effects requires intense future studies, it is obvious that *tag-63* knockout (as well as knockdown) negatively affects anterograde movements of UNC-104. We also plotted kymograph traces of motors and synaptic vesicles (marked by SNB-1) (Figure 6C) into bullet diagrams (Figure 7A-D) and from these diagrams, it is apparent that UNC-104 motors move longer distances in a shorter period of time in mutants (as opposed to wild types), specifically obvious in retrograde directions (Figure 7A+B). Indeed, single event run lengths (for definition see Suppl. Figure S9D) are increased in retrograde directions in TAG-63 KO worms (Figure 7G). From bullet diagrams, it is also evident that net run lengths (defined by the distance between the starting point and the end point of an observed particle run, Suppl. Figure S9D) are increased (Figure 7B). Likewise, quantification of these data revealed a significant increase in average net run-lengths in retrograde directions for UNC-104 in *ok471* mutant animals (Figure 7E). We also measured an increased amount of single retrograde event run lengths (Figure 7G, and for definition of “event run length” refer to Suppl. Figure S9D). Increased retrograde speeds are furthermore obvious when examining the representative kymographs in Figure 6C. Increased retrograde speeds observed for SNB-1 particles are consistent with the determined reduction in anterograde event run lengths (Figure 7G). Also note that upon administration of acrylamide (600 mg/ml) both anterograde and retrograde axonal transport was significantly inhibited (Figure 6A+B; also see kymograph in Figure 6C) consistent with previous studies on the role of NFs on axonal transport ^47-49^.

These measured effects reveal visible consequences on axonal transport: by observing UNC-104 particle distribution in ALM neurons (Figure 8A+C), it is evident that decelerated UNC-104 in *ok471* mutant worms lead to measurable accumulation (clustering) of motors along axons (Figure 8F). Vice versa, increased retrograde velocities and decreased anterograde single event run lengths, as seen for SNB-1 in *ok471* mutant worms, lead to visibly shorter travel distances (Figure 8B+D and E). These effects of TAG-63 KO on UNC-104 (reduced anterograde velocities and increased motor clustering) and SNB-1 (increased retrograde velocities and reduced anterograde travel distances) are also directly visible in time-lapse image sequences presented in Suppl. Video S1. Note that mRNA expression levels of unc-104 do not vary between wild type animals and *tag-63* mutants (Suppl. Figure S9C).

### Abnormal targeting of synaptic vesicle precursor (SNB-1) in *tag-63(ok471)* mutants

The HSN neuron is widely used as a model to study synapse function and development. Figure 8G shows a schematic representation of a HSN neuron with synapses marked in purple that form along vulval muscles. Because we determined significantly diminished anterograde run-length of SNB-1 in *tag-63* mutants, it would be interesting to understand whether SNB-1 precursor targeting to synapses would be also compromised. Indeed, SNB-1 largely accumulates in HSN cell bodies of *tag-63* mutants (but not in wild types) obviously unable to reach HSN synapses via fast axonal transport (Figure 8H+I). At the same time, SNB-1 *en passant* puncta number (along sublateral neurons) are reduced in *tag-63* mutants consistent with aforementioned observed SNB-1 retention phenotypes in somas of HSN neurons in these mutants (Figure 8J+K). Critically, two other cargoes in the fast axonal transport system RAB-3 (known to be transported by kinesin-3) and mitochondria (known to be transported by kinesin-1) do not display any visible transport defects in HSN neurons (Suppl. Figure S10A-D).

### Interactions between UNC-104/TAG-63 and SNB-1/TAG-63

As a first approach to understand whether or not functional interactions between UNC-104/TAG-63 and SNB-1/TAG-63 may exist we carried out colocalization experiments and used intensity correlation quotient (ICQ) for quantification. As a result, ICQ values in worms either co-expressing *unc-104::mrfp* and *tag-63::gfp* or *snb-1::mrfp* and *tag-63::gfp* were all positive (Figure 9A-C and Suppl. Figure S11A+C) pointing to interdependent colocalization (for ICQ value definition refer to Materials & Methods). Moreover, exposing worms to acrylamide resulted in significant reduction of colocalization between UNC-104 and TAG-63, as indicated by the Pearson’s correlation coefficient (Figure S11B+D). To further understand whether or not these proteins are indeed close enough to form functional interaction complexes we then carried out bimolecular fluorescence complementation (BiFC) assays. Here, the assayed protein pairs are fused to a non-fluorescent YFP hybrid (either VC or VN) and concomitant YFP signals (due to successful complementation of hybrids) would indicate that the two test proteins are less than 7 nm apart. UNC-104/UNC-104 BiFC was used as positive control and UNC-104/empty BiFC was used as negative control (Figure 9D). Notably, UNC-104/TAG-63 and SNB-1/TAG-63 both exhibited positive BiFC signals indicating the likelihood for physical interactions (Figure 9D). Note that we also observed colocalization of UNC-104 and SNB-1 in sublateral nerve cords and that colocalization was significantly reduced in *tag-63* mutants (Figure 9E+F).

## DISCUSSION

To dissect the underlying molecular mechanisms of human neurological diseases, model organisms such as *Zebrafish, Drosophila*, and *C. elegans* are frequently employed ^50,51^. Specifically *C. elegans* has been increasingly used to study neurodegenerative diseases such as tauopathies, ALS, AD, PD and CMT^5,52,53^. While many “disease genes” can be conveniently studied in nematodes, including the tau homolog *ptl-1*, APP-related gene *apl-1*, and presenilin homolog *sel-12*, no NF homolog has been identified so far. Notably, a plethora of brain diseases are associated with genetic defects in NF genes leading to disintegrated NF networks, often visible in pathological NF aggregates ^2,15,54^. Critically, phenotyping the *tag-63* mutant worm unraveled reduced responses to soft and harsh touch, and these abnormal behaviors can be related to mechanosensory neuron defects (Figure 2). The very obvious levamisole resistance phenotype (Figure 2C) may point to downregulation of nicotinic acetylcholine receptors at NMJ’s ^55^. Furthermore, since egg-laying is regulated by HSN neurons^56^, the observed egg retention phenotype (Figure 2A+B) is consistent with determined synaptic vesicle accumulations in HSN somas together with reduced number of vesicles at HSN synapses (Figure 8G-K). With such obvious mechanosensory neuron and HSN neurons defects, together with nicotinic acetylcholine downregulation, the *ok471* worm may serve well as a disease model to study TAG-63/neurofilament-based diseases. Specifically, it would be interesting to identify genes that would suppress *tag-63* phenotypes.

From this study it is obvious that TAG-63 on one hand reveals similarities to NFH (orthologies, Figure S2; sequence homologies, Figure S(1+4+5)) while its structural features resembles rather that of NFL (lack of KSP repeats, Figure S3; protein size, Figure 5A+B; ability to form filaments via homopolymerization, Figure 5) [note that the WormBase BLAST search shown in Figure S1 can be only reproduced with database #WS230 because later versions use different algorithms delivering different results]. TAG-63 is expressed in a wide subset of neurons, including ventral nerve cord (comprising most motor neuron soma), sublateral neurons, mechanosensory neurons, CANL/CANR neuron, tail neurons as well as amphid neurons (sensory neurons) (Figure 3, Suppl. Figure S7). Western blots detect TAG-63 protein using anti-NEFH antibodies and the band disappears in KO worms (Figure 1E+F). Anti-NFH antibodies also stain *C. elegans* neurons and this staining is reduced in both TAG-63 KO worms and in TAG-63 RNAi worms (Figure 4A-E and S8A+B). Primary cultured *C. elegans* neurons are sensitive to acrylamide (ACR) and tricresyl phosphate (TCP), and treatment with these substances induce axonal development defects (Suppl. Figure S9A+B), likely as a result of a disrupted NF network (known to mechanically stabilize delicate axons) ^37-39^. Indeed, on the molecular level we can see that *in vitro* reconstituted TAG-63 filaments disintegrate upon exposure to acrylamide (Figure 5G) (similar as observed for neurofilaments ^42^). Notably, the toxic effects of acrylamide have been discussed as a cause for neurofilament-related axonopathies ^57^.

From TEM images it is obvious that purified TAG-63 is able to form polymeric structures including bundles with occasionally visible 10 nm filaments (Figure 5). Note that homopolymerization capacity observed for TAG-63 is also a characteristic feature of human NFL ^15,16^. Urea is known to be a critical denaturant to solubilize NF proteins, and after urea solubilization, TAG-63 indeed forms globular structures of sizes ranging from 120 nm to 180 nm (Figure 5C), which very much resembles those seen for denaturized NF proteins ^18^. Further, as reported by Krishnan *et al.* ^18^, these globular structures are able to convert into 10 nm filaments upon prolonged dialysis against Tris-buffer, a phenomenon which can be also seen for TAG-63 (Figure 5). Note that IFs are temperature sensitive and form stable polymorphic structures depending on the incubation temperature ^58^. Indeed, we identified two variants of TAG-63 structures depending on the temperature: thick aggregates (Figure 5D) can be detected at higher temperatures (about 37°C), and fibrous bundles with emanating single filaments (Figure 5E+F) of 10 nm size at lower preparation temperatures (22°C).

Because NFs often run in parallel and are closely associated with MT networks ^40^, we speculated that fast axonal transport may be modulated in TAG-63 knockout (KO) animals. Indeed, anterograde axonal transport is significantly impaired in *tag-63* mutants and knock-down worms (Figure 6+8), likely due to slower velocities of the transporter kinesin-3 UNC-104 (Figures 6+7). It has also been reported that acrylamide treatment may directly or indirectly affect anterograde axonal transport by penetrating the nerve barriers ^47^. Moreover, a single acrylamide injection caused degeneration of motor neurons in hens leading to deficits in retrograde axonal transport ^49^. Consistent with these studies, we determined that acrylamide administration to worms adversely affects the anterograde and retrograde axonal transport of UNC-104 (Figure 6A+B). Interactions between NFs and organelles as well as molecular motors have been frequently reported. For example, it has been demonstrated that NF and synaptic vesicles formation are co-delayed in a rat model of gastroschisis ^43^, and NF depletion co-occurs with synaptic vesicle reduction after exposing rodents to the neurotoxicant 3,3’-imino-dipropionitrile (IDPN) ^45^. Further, it has been reported that NFs regulate NF-kinesin and NF-NF associations *in vivo* ^59^. Likewise, the roles of kinesin-1 and dynein on NF transport were thoroughly characterized ^25^. These reports from others, together with our observations (Figure 6-9), point to a functional role of TAG-63 (a NF-like protein) on axonal transport, and we suggest a model in which the lack of NFs or TAG-63 may lead to impaired transport of SNB-1 tagged synaptic vesicles (Figure 10, and discussion below). Interestingly, two other analyzed axonal cargoes such as RAB-3 vesicles or mitochondria remain unaffected by TAG-63 KO (Figure S10) pointing to the specificity of observed effects for SNB-1 vesicles. Also note that unc-104 mRNA expression levels are unaffected in *tag-63* mutant worms (Suppl. Figure S9C) demonstrating that observed effects are likely not based on transcriptional changes but on factors that directly regulated UNC-104 motors. Besides the indication that TAG-63 may directly affect UNC-104 powered synaptic vesicle transport, studies have also shown that NFs attenuate tau-mediated axonal transport defects and neurite retraction ^44^. Here, tau overexpression leads to excessive tau-kinesin associations, which in turn deactivates the anterograde- and activates the retrograde transport-machinery, whereas NF co-expression was able to attenuate these transport defects. These findings may partially explain the observed decrease in anterograde velocities of kinesin-3 motors together with increased retrograde velocities of SNB-1 particles in TAG-63 KO worms (Figures 6+7). Moreover, our group has shown previously that PTL-1/tau associates with UNC-104 and depletion of PTL-1 negatively affects retrograde axonal transport (but not anterograde axonal transport) ^31^.

**Figure 10:**
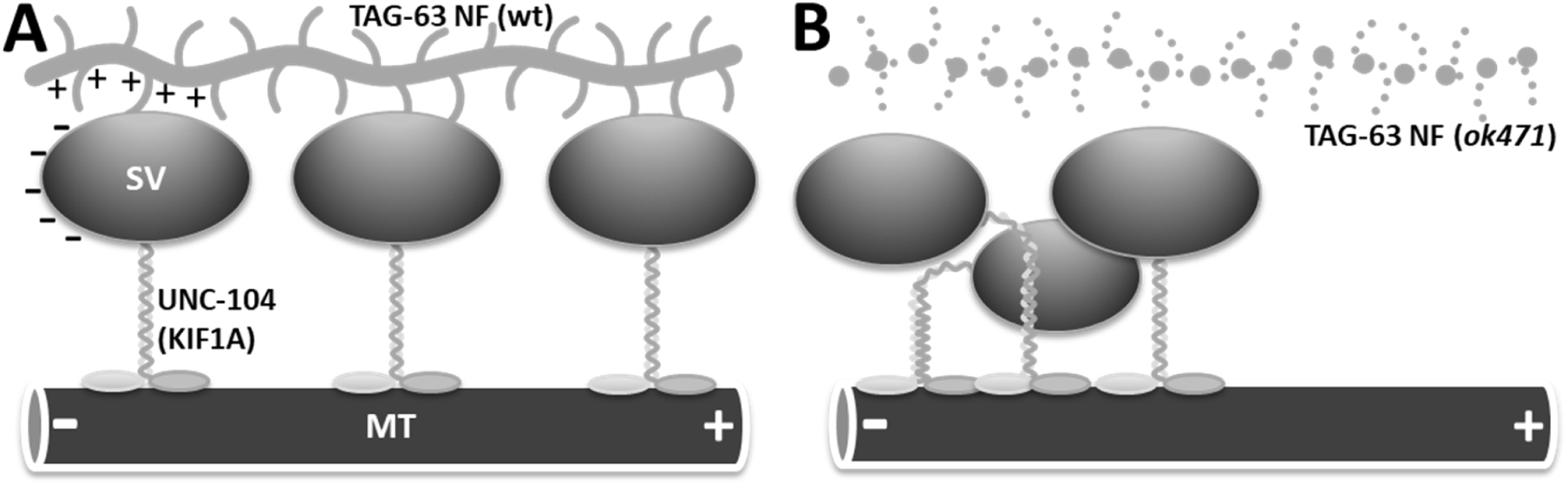
Model depicting the effect of *tag-63* on axonal transport. (A) In wild type nematodes, synaptic vesicles (SV, marked by SNB-1::mRFP) are transported by UNC-104 into anterograde directions. Not shown is a proposed effect by the retrograde motor dynein, which, when activated, antagonizes this movement by pulling the vesicle into opposing directions (MTs minus ends). While the motor is tightly bound to the MT, the (comparatively) large vesicle also faces certain support by dynamic and transient (likely weak electrostatic) interactions with closely parallel running TAG-63 filaments. This support would facilitate the motor’s processivity expectedly due to increased attachment rates (positively affecting its duty cycles). (B) In *tag-63* mutants, stabilization of the cargo is lost, the interaction between UNC-104 and SNB-1 is reduced, and thus motor processivity is negatively affected. As a result, UNC-104 motor motility and synaptic vesicle transport are compromised (see Discussion for details).

Besides NFs, vimentin has been reported to regulate organelle transport. Specifically, mitochondrial morphology ^60^, motility ^61^, and membrane potential ^24,62^ are all affected by vimentin. Furthermore, interactions between vimentin and molecular motors have been reported, particularly, vimentin interacts with kinesins to establish cross-linked cytoskeletal networks ^63-65^. Also, vimentin interacts with dynein during Ndel1-mediated neurite outgrowth ^66^, and during the transport of injury signalling molecules ^67^ as well as during nuclear rotation ^68^. Based on these studies, a general role of IFs on regulating intracellular transport mechanisms may be assumed. In our model (Figure 10), NFs (closely associated with the microtubule-based transport machinery) stabilize the motor-cargo complex by providing weak and transient (surface charge-based) interaction points. Therefore, not only the motor is properly bound to a supporting track (via microtubules), but also the vesicular cargo (via neurofilaments). Since the cargo itself is often of magnitudes larger than the motor, it would be plausible that a stabilized cargo would be more efficiently transported than a “free-floating” cargo. Hence, we propose that during their travels along axons, organelles, such as synaptic vesicles, transiently dock and undock onto NFs based on weak electrostatic interactions. Indeed, the synaptic vesicle surface is negatively charged ^69^, and NF tails (side arms) are positively charged ^13,70^, and it is noteworthy that “charge regulation” plays a major role in protein-protein and protein-membrane interactions ^71^. We calculated a net isoelectric point of 8.31 for TAG-63 with, interestingly, an elevated isoelectric point of 9.31 at the C-terminus, suggesting highly positive charged tails (isoelectric.ovh.org) similar to NF side-arms. Also, it has been reported that NF-mitochondria interactions depend on mitochondrial membrane potential as well as the extent of NF side arm phosphorylation ^14^. Another important feature of our model is that NF-mediated cargo stabilization would positively affect the activation of UNC-104 because it is thought that UNC-104 remains inactive (via intramolecular folding) if not bound to a proper cargo ^72-74^. Also, motors are often activated when experiencing mechanical load caused by the cargo ^75-77^. Nevertheless, besides a direct interaction between NFs and cargo (such as inactivated dynein bound to a synaptic vesicle), we also may consider that the lack of TAG-63 would modulate MT dynamics, leading to observed effects as shown in Figure 6-8. Indeed, it has been shown that NFs inhibit the polymerization of MTs by binding to both MAPs and unassembled tubulin ^78,79^, and that MTs are present in abnormal numbers in NF-deficient axons ^78^.

## MATERIALS & METHODS

### C. elegans maintenance and used strains

Worms were maintained at 20°C on NGM agar plates seeded with OP50 *E. coli* according to standard methods ^80^. To clone a *Ptag-63::gfp* transcriptional fusion, we amplified a 3 kb promoter using forward primer CTGCAGGCGTTGTTGAAAATCTCC and reverse primer GTCGAC-CTGAAAAAATGTAAATAATTCAA-TAAACAAGAAAAGTTTTA, and then subcloned the amplicon into the pPD95.77 promoterless GFP vector (Addgene, MA, USA). The plasmid was then coinjected along with *Punc-104::unc-104::mrfp* (both at 100 ng/μl) into N_2_ worms ^31,46^ to receive OIW 52 *N*_*2*_;*nthEx52*[*Ptag63::gfp*;*Punc-104::unc-104::mrfp*].

To clone a *Ptag-63::tag-63::gfp* translational fusion, we designed primers ACATTTTTTCAGCCCATGCCGCG-TATCAAGCAA (forward) and CCTTT-GGCCAATCCCTTACGTCGAACACCA-TAG (reverse) for *tag-63* cDNA amplification and then directly cloned the PCR amplicon into pPD95.77 *Ptag-63::gfp* construct using the FuseIn™ cloning technique. The plasmid was coinjected along with *Punc-104::unc-104::mrfp* (both at 100 ng/μl) into *tag-63(ok471)* mutant worms to generate OIW 53 *tag-63(ok471);nthEx53 [Ptag63::tag-63::gfp;Punc-104::unc-104::mrfp]* rescue strain.

To clone a *Ptag-63::tag-63::VN173* translational fusion, we first cloned Ptag-63 promoter into pPD95.77 promoterless VN173 vector to create a transcriptional fusion. *pPD95.77 VN173* was cloned by replacing GFP in the expression vector with VN173 using the forward primer AAAAAAGGTACCGG-TAGAAAAAATGGTGAG-CAAGGGCGAGGA and AAAAAA-GAATTCTACGAATGCTCGATGTT-GTGGCGGATCTTGAAGTT reverse primer. Next, Ptag-63 promoter was amplified using AAAAAACCTGCAGGGCGTTGTT-GAAAATCTCC forward primer and AAAAAACCCGGGCTGAAAAAATGTAAATAATTCAATAAAC reverse primer and then the 3 kb promoter was subcloned into pPD95.77 promoterless VN173 vector. Further, to create *Ptag-63::tag-63::VN173*, we amplified *tag-63* gene from cDNA using ACATTTTTTCAGCCCATGCCGCG-TATCAAGCAA forward and CCTTT-GGCCAATCCCTTACGTCGAACACCA-TAG reverse primers and subcloned the gene into *pPD95.77 Ptag-63::VN173* vector using the FuseIn™ cloning technique. This plasmid was coinjected along with either *Punc-104::unc-104::VC155* or *Punc-104::snb-1::VC155* plasmid shown in earlier publications ^46^ (both at 75 ng/μl) into *N*_*2*_ worms to generate OIW 54 *N*_*2*_;*nthEx54 [Ptag63::tag-63::VN173;Punc-104::unc-104::VC155]* and OIW 55 *N*_*2*_;*nthEx55 [Ptag63::tag-63::VN173;Punc-104::snb-1::VC155]* strains respectively.

To clone TAG-63 L4440, the cDNA sequence between the start and stop codon of *tag-63* was cloned using primers AAAAGGTACCCCGCGTATCAAGCAAGC (forward) and AAAACTGCAGACGTCGAACACCATAGCC (reverse) into the L4440 vector. The obtained clones were transformed into HT115 competent cells for RNAi feeding assays. Similarly, *snb-1* genomic DNA was cloned into an already existing *pPD95.77 Punc-104::mrfp* vector using the primers CTCTAGAGGATCCCCATGGACGCTCAAGGAGATGCC (forward) and CCTTT-GGCCAATCCCTTTTTTCCTCCAGCCCA-TAAAACGATGAT (reverse). OIW 56 *N*_*2*_; *nthEx56[Punc-104::snb-1::mrfp; Ptag-63::tag-63::gfp]* was generated by injecting SNB-1::mRFP and TAG-63::GFP at a concentration of 80 ng/µl and 120 ng/µl, respectively.

To visualize pan-neuronal UNC-104::mRFP expression in *tag-63* knockout (KO) strains, we crossed the existing strain *Punc-104::unc-104::mrfp (e1265)* with VC275 *tag-63*(*ok471*) (which was three times outcrossed after receiving it from the *Caenorhabditis* Genetics Center, CGC) to receive strain OIW 57 *tag-63*(*ok471*);*nthEx57*[*Punc-104::unc-104::mrfp*]. To visualize synaptic vesicles (identified by SNB-1::mRFP expression), an existing strain co-expressing SNB-1::mRFP and UNC-104::GFP ^81^ was crossed with the VC275 *tag-63*(*ok471*) strain to receive OIW 58 *tag-63*(*ok471*);*nthEx58*[*Punc-86::snb-1::mrfp*;*Punc-104::unc-104::gfp*].

To amplify *rab-3* gene from cDNA, we used AAAAGCTAGCATGGCGGCTGG-CG forward and AAAACCATGGTTAG-CAATTGCATTGCTGTT reverse primers respectively. After NheI and SalI digestion, *rab-3* was ligated to the *Punc-104::mcherry::GW* expression vector to create *Punc-104::mcherry::rab-3* construct. This plasmid was injected at a concentration of 80 ng/μl into *N*_*2*_ worms to generate OIW 59 *N*_*2*_;*nthEx59 [Punc-104::mcherry::rab-3]* and OIW 60 *tag-63(ok471);nthEx60[Punc-104::mcherry::rab-3]* strains respectively.

To generate *Punc-104::mito::gfp* plasmid construct, 5 kb upstream of *unc-104* gene was amplified using the following primer set: AAAAAAGCTTGTCATTTTTT-GTTTTTATTTATTTTCCAAAAAAAC-CTATTTTTTTGTTT (forward) and AAAAGTCGACCTGAAATT-GTTTTAATTAATTCAAGAATTCGTTCG-TAGAAATTG (reverse) primers, and subcloned into pPD96.32 encoding mitochondrial targeting sequence and GFP. Thus obtained plasmid was microinjected at the concentration of 75 ng/µl to generate OIW 61 *N*_*2*_;*nthEx61[Punc-104::mito::gfp] and* OIW 62 *tag-63(ok471);nthEx62 [Punc-104::mito::gfp]* strains respectively.

### Behavioral assays

Egg retention assay was carried out as described by Gardner *et al.* ^82^. In brief, L4 worms were spread on an NGM agar plate with OP50 and incubated at 20°C for 40 hours. After 40 hours, the number of eggs inside the worm was counted using bright-field microscopy. For Figure 2B, laid eggs were counted each day. Quadrant chemotaxis assay was carried out as described previously ^83^ employing benzaldehyde (0.5% mixed with equal volume of sodium azide 0.5 M) as a chemoattractant. Chemotaxis index was calculated for three independent trials each with 19∼25 worms. Thrashing frequency was determined as the worms’ body bends per minute in M9 buffer, pharyngeal pumping (analyzed on OP50 NGM plates) was counted manually under a stereo microscope ^84^, and touch response was analyzed as described previously ^85^. For levamisole and aldicarb paralysis assays NGM plates were prepared with 0.2 mM levamisole (Sigma #T1512) and 1 mM aldicarb (Sigma, #33386). About 25 worms were added to these plates and scored for paralysis every 30 minutes for 8 hours as described by Butler *et al.* 2018 ^86^. Worms lost during the assay time were excluded from the analysis. Experiments were repeated three times.

### Real-time PCR

To evaluate tag-63 and unc-104 mRNA levels in *ok471* mutants (Figure 1D), we performed real-time PCR assays employing ABI StepOne Plus Real time PCR system in conjunction with the ABI Power SYBR green PCR master mix. The following primers were used: ATGCCGCGTATCAAG (forward) and TTT-GTTCGAGTCGAGC (reverse) that encompass the 1st and 2nd exon (178 bp) for *tag-63*. For quantifying unc-104 mRNA expression levels the following primer set was used: GAAGGAAATAAAGCGAGAAC (forward) and CTGCCAAATCAACCAAAG (reverse) (Figure S9C). Messenger RNA levels of both *tag-63* and *unc-104* were normalized to *cdc-42* internal control with CTGCTGGACAGGAAGATTACG (forward) and TCG-GACATTCTCGAATGAAG (reverse) primers respectively.

### Western blotting from whole worm lysates, neuronal cell isolation and immunohistochemistry

To perform Western blotting from whole worm lysates (Figure 1E+F), we employed a previously published protocol ^87^, using 100 µg sample protein and membrane blocking by milk powder for 1-1.5 hours at room temperature (RT). Primary anti-NEFH antibody (mouse, monoclonal, #WH0004744M1, Sigma) was incubated at 4°C for 14 hours at 1/500 dilution. The secondary anti-mouse antibody (#sc-2005, Santa Cruz) was incubated at RT for 2 hours with 1/500 dilutions. For treatment of cultured neurons with acrylamide (5 mM and 30 mM) and TCP (1 µg/ml) (Suppl. Figure S9A+B), we isolated primary neurons from *C. elegans* embryos based on protocols by Christensen *et al.* ^88^ and Strange *et al.* ^89^. Isolated neurons are maintained in Leibowitz’s L-15 medium with 10% fetal bovine serum, 100 U/ml penicillin, 100 μg/ml neomycin, and 100 μg/ml streptomycin. Nematode cells can be stored at RT without the need to adjust the CO_2_ atmosphere. For the treatment of worms with acrylamide, we added 600 mg/ml acrylamide to NGM plates as described previously ^90^ (Figure 6A+B).Whole mount immunostaining was carried out based on the tube fixation method published previously ^91^. Here, worms were fixed with 4% formaldehyde for 30 minutes, followed by a freeze-thaw cycle to crack open the epidermis and again a second round of fixation for 30 minutes. After β-mercaptoethanol (3 hours at RT) and collagenase treatment (24 hours at 37°C), worms were blocked with donkey serum and stained with primary and secondary antibodies. Primary antibody was either anti-NEFH antibody (produced in mouse, monoclonal, #WH0004744M1, Sigma) or anti-NFH 200 (produced in mouse, monoclonal, #N5389, Sigma). Secondary antibody was either goat anti-mouse IgG H&L Alexa Fluor 647 (Abcam, ab150115) or anti-mouse IgG (H+L), F(ab’)2 fragment, CF 488A (SAB4600388-125UL, Sigma). Primary antibody dilution was 1:50 and the secondary antibody dilution was either 1:250 (Figure 4A+B, S8C) or 2.5:1000 (Figure 4D).

### RNAi feeding assay

NGM plates containing ampicillin (25 μg/ml) and 1 mM IPTG were inoculated with HT115 *E. coli* strain carrying the appropriate dsRNA clone, and grown overnight. 15-20 worms were transferred to RNAi feeding plates and incubated at 20°C. Worms were then transferred to new RNAi feeding plates every 24 hours and the F1 progeny was used for analysis after at young adult stage. To analyze knockdown efficiency, we performed *tag-63* RNAi on TAG-63::GFP expressing worms (Figure S8A+B). For quantifying immunostaining of NFH 200 (#N5389, Sigma) (Figure 4D+E), HT115 bacteria transformed with L4440 TAG-63 ORF were fed to age synchronized worms. Worms were transferred every 24 hours to fresh RNAi plates. As control, worms were fed with HT115 bacteria containing L4440 empty vector.

### Recombinant protein expression, solubilization and purification

Tag-63 cDNA was cloned into PET expression vector pET-23a (+) that carries an N-terminal T7 tag sequence along with a C-terminal His tag sequence. The cDNA was directly cloned into the PET vector using GGATCCGAATTCGAGATGCCGCG-TATCAAGCAAGC (forward) and AGCTT-GTCGACGGAGAAACGTCGAACACCA-TAGC (reverse) primer by FuseIn™cloning technique. *E. coli* BL21 (DE3) was used as a host for the production of recombinant proteins. Plasmids carrying the *tag-63* gene were transformed into *E. coli*. Colonies were then cultured in LB containing 50 µg/ml ampicillin followed by incubation at 37°C until the OD_600_ reached a value between 0.6-0.8. The culture was then induced with IPTG (0.1 mM) followed by incubation at 16°C for 16 hours. Cells were harvested by centrifugation at 4000 rpm at 4°C for 15 minutes and then suspended into 1/10 volume of Tris-buffer followed by sonication on ice. Cell debris and supernatant were separated by centrifugation at 13,000 rpm for 30 minutes at 4°C. The pellet was resuspended in 0.5 ml urea lysis buffer containing 8 M urea and incubated at 30°C for 30 minutes. The solubilized supernatant was collected and analyzed by 8% SDS-PAGE. Selected colonies with optimized induction were further employed in large scale expressions. 3 ml nickel sepharose was poured in a 1.5 × 10 cm column followed by washing with 5 folds column volumes of ddH_2_O and equilibrated with 30 ml binding buffer. Protein samples were then loaded into the columns and washed with 50 ml binding buffer followed by 50 ml wash buffer (containing 50 mM imidazole). The sample was then eluted with elution buffer (containing 300 mM imidazole). Lastly, purified proteins were dialyzed against 20 mM Tris-buffer at 4°C for 36 hours and analyzed by 8% SDS-PAGE using rabbit polyclonal 6X His Tag antibody (GTX115045), anti-NEFH antibody (produced in mouse, monoclonal, #WH0004744M1, Sigma), and anti-NFH 200 (produced in mouse, monoclonal, #N5389, Sigma) at a dilution of 1/1000 and a rabbit IgG (HRP) 2° antibody (GTX213110) and antimouse 2° antibody (#sc-2005, Santa Cruz) at a dilution of 1/1250 (Figure 5B).

### Rhodamine staining and acrylamide treatment of reconstituted TAG-63 filaments

NHS-Rhodamine (10 mg/ml, Thermo Scientific Product no 46406) was dissolved in DMSO and mixed with purified TAG-63 protein. This dye-protein sample was then incubated at room temperature for 1 hour followed by removal of the excess dye by a dye-removal column. Lastly, the rhodamine-labeled protein was stored at 4°C protected from light before imaging with an epi-fluorescence microscope (Figure 5G (a)). Rhodamine-stained TAG-63 filaments were also treated with acrylamide in varying concentrations (0.1 mM - 0.5 mM) following 16 hours of incubation on a rotary shaker (Figure 5G (b-f)).

### TEM analysis

Sample droplets (10 µg) were placed onto carbon coated copper grids (400 mesh) and excess supernatant fluid was removed using filter paper. A droplet of contrasting agent (2% uranyl acetate) was added, blotted and airdried. Imaging was performed using a Hitachi HT7700 Transmission Electron Microscope at 100 kV (Figure 5C-F).

### Worm imaging and motility analysis

For microscopic observations, worms were immobilized on 2% agarose-coated cover slides, and no anesthetics (e.g., levamisole) were used (reported to affect motor motility ^92^). An inverted fluorescence microscope (Observer.Z1, Zeiss) was used for imaging worms as shown in Suppl. Figure S7 and a Zeiss LSM780 confocal laser scanning microscope with 100X objective was employed for imaging worms in Figures 3, 4, 8H, 8J, and 9 as well as in Suppl. Figure S8, S10 and S11. For motility analysis and imaging of axonal clustering (Figure 6-8(A-D)), we employed an Olympus IX81 microscope with a DSU Nipkow spinning disk unit connected to an Andor iXon DV897 EMCCD camera for high-resolution and highspeed time-lapse imaging at 3-4 frames per second. To convert recorded time-lapse sequences into kymographs, we used the imaging analysis software NIH ImageJ (NIH, http://rsb.info.nih.gov/ij/). The “straighten plugin” was used to straighten curved axons, and after drawing a line over the axon, the plugin “reslice stack function” was executed to generate kymographs. In kymographs, static particles appear as vertical lines, whereas the slope of a moving particle corresponds to the velocity (speed) of the particle (Suppl. Figure S9D). A pause is defined if motors move less than 0.07 μm/s, and each calculated velocity event does not contain any pauses. On average, we analyzed approximately 600 moving events per experiment from 15 to 25 individual worms. A moving event is defined as a single motility occurrence typically right after a pause or a reversal, and an event ends when the motor again pauses or reverses (Suppl. Figure S9D). Motor cluster analysis (Figure 8F) was carried out using ImageJ’s “area” tool and the “particle analyze” plugin. To measure travel distances of particles in (straightened) neurons (Figure 8E), we used ImageJ’s “line tool” to draw a line from the proximal axon hillock to the identified distal particles. Similarly, we measured the lengths of axons in cell cultures grown with or without the particular drug (Suppl. Figure S9B). Intensity Correlation Quotient (ICQ) was calculated by selecting the region of interest using the “polygonal selection tool” in ImageJ. After background fluorescence intensity subtraction, the plugin ‘Intensity Correlation Analysis’ was executed to generate either ICQ or Pearson’s correlation coefficient (PCC) values (Figure 9C+F and S11D). ICQ values range from −0.5 to 0.5, and values close to 0.5 stand for interdependent expression of two fluorophores, values close to −0.5 stand for segregated expression, and values around 0 imply random expression. Pearson’s correlation coefficient values vary from +1 to −1 and +1 stands for complete correlation, 0 for no correlation and −1 for complete anti-correlation ^93^. For fluorescence intensity quantification the following formula was used: Integrated density of selected region – (area of selected region * mean fluorescence of the background) ^94^. Parameters such as mean grey value of background, area of selected region and integrated density of selected region were measured using ImageJ (Figure 4C+E, 8I, S8B+D and S10B+D). For figure 8I and S10B+D we have calculated the percentage of fluorescence intensity in soma and synapse by normalizing their individual values with the sum of fluorescence intensity from soma and synapse which is 100%. Line scans were generated using ImageJ’s “plot profile” plugin after straightening the neurite (Figure S8D).

## ACKNOWLEDGEMENTS

We thank Dr. Nai-Wen Tien for assisting with the acrylamide and TCP experiments on cell cultures. We also like to acknowledge Prof. Margaret Dah-Tsyr Chang and her lab members Yi-En Chen and Tse-Kai-fu (Institute of Molecular and Cellular Biology, NTHU, Taiwan) for providing various chemicals and aiding us in TAG-63 protein expression and purification experiments. We thank the *C. elegans* Core Facility (CECF) Taiwan (funded by the Ministry of Science and Technology, MOST) for providing microinjection setups and worm observation systems. We acknowledge MOST grants 103-2311-B-007 -004 -MY3 and 101-2311-B-007-002- to OIW and support by the Brain Research Center of National Tsing Hua University under the Higher Education Sprout Project funded by the Ministry of Science and Technology and Ministry of Education in Taiwan.

This paper is dedicated to our friend and collaborator Dr. Jean-François Leterrier (Université de Poitiers, France) who contributed enormously to the field of neurofilament research and passed away much too soon in 2017.

## Author Contributions

PB performed experiments, analyzed the data and wrote the paper. MMS performed experiments, analyzed the data and wrote the paper. DW performed experiments and analyzed the data, OB performed experiments and CWC analyzed the data. OIW designed experiments, secured funding and wrote the paper.

## SUPPLEMENTARY MATERIAL

### SUPPLEMENTARY FIGURE LEGENDS

**Suppl. Figure S1:**
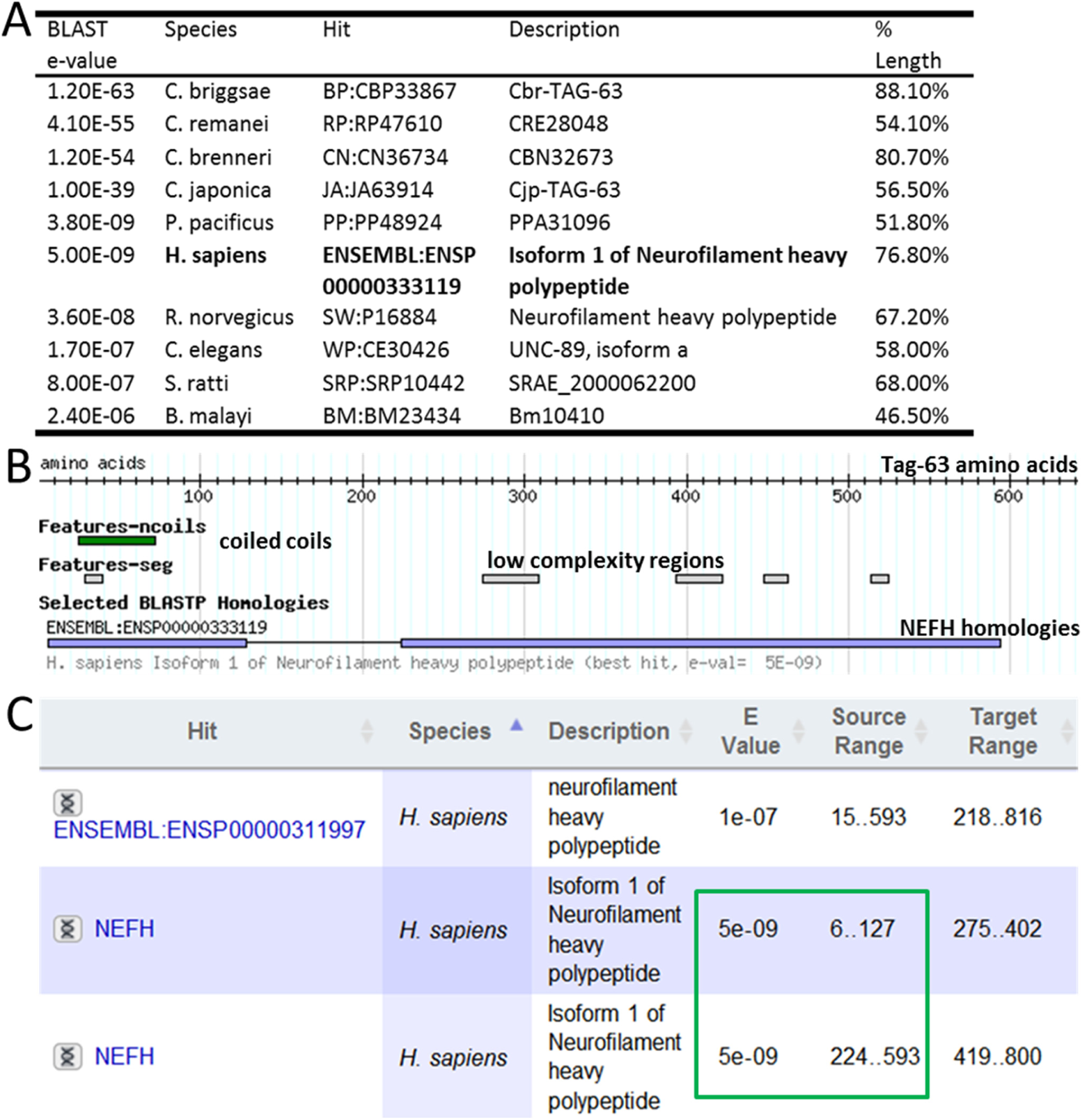
WormBase BLAST (database version #WS230) using TAG-63 as a query. (A) Table with results from WormBase TAG-63 BLAST using database version #WS230 (wormbase.org). (B) WormBase TAG-63 protein features with marked (blue) NEFH homologies, and corresponding E values (framed in green) shown in (C). Note that to reproduce the data shown in this Suppl. Figure, WormBase database #WS230 must be used since later database versions employ different algorithms leading to different results (WormBase BLAST version #WS230 employs complex AB-BLAST algorithms (advbiocomp.com/blast.html) aggregating various databases such as ENSEMBL, SWISS-PROT and TrEMBL (wiki.wormbase.org), likely enabling for more in-depth searches).

**Suppl. Figure S2:**
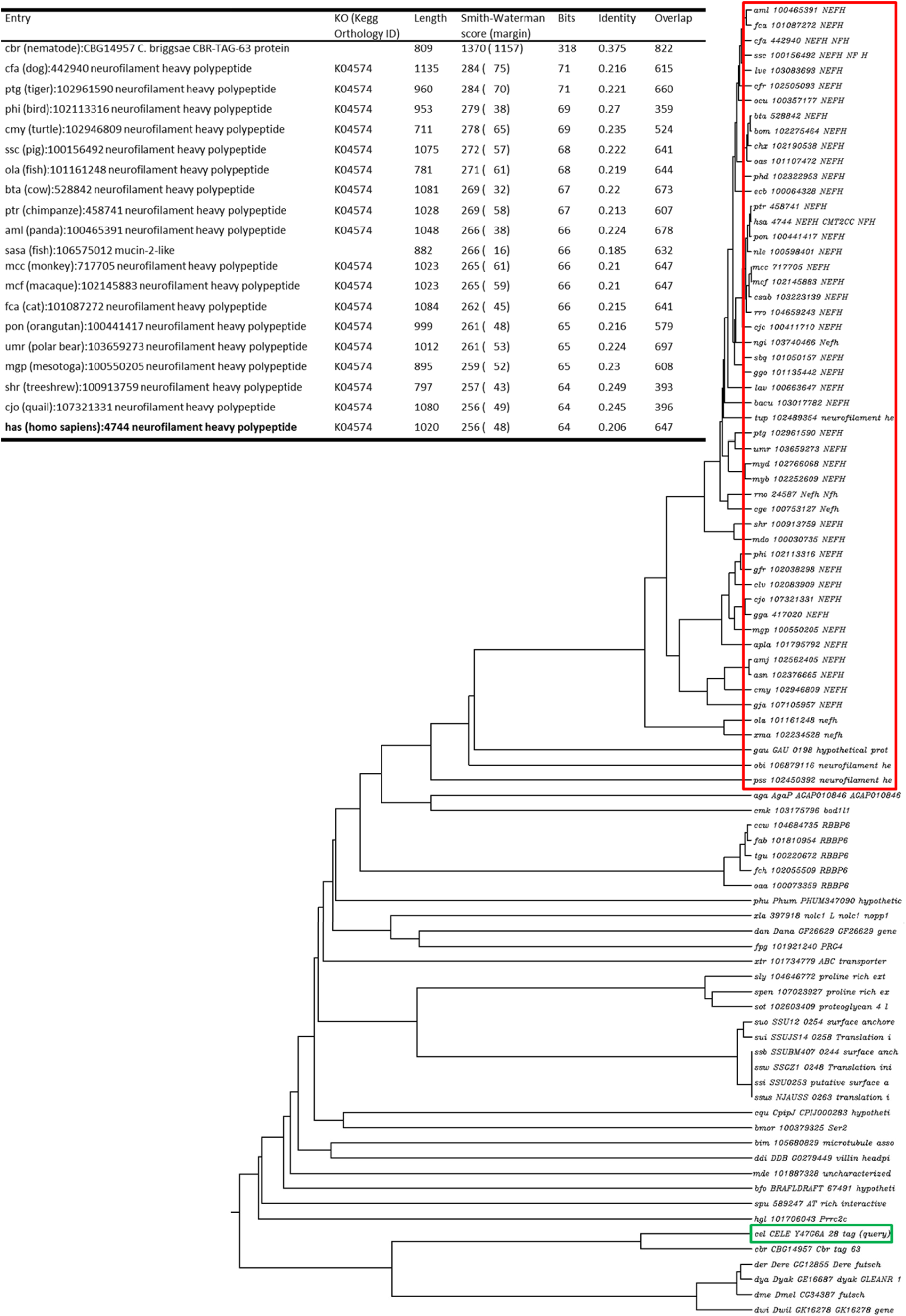
KEGG BLAST results and phylogenetic tree. KEGG BLAST with CELE_Y47G6A.28 as query reveals various NEFH orthologs (kegg.jp). KEGG rooted phylogenetic tree (dendrogram) displays a large cluster of various NEFH encoding genes (framed in red) that have emerged from a common ancestor from which TAG-63 also emerges (framed in green).

**Suppl. Figure S3:**
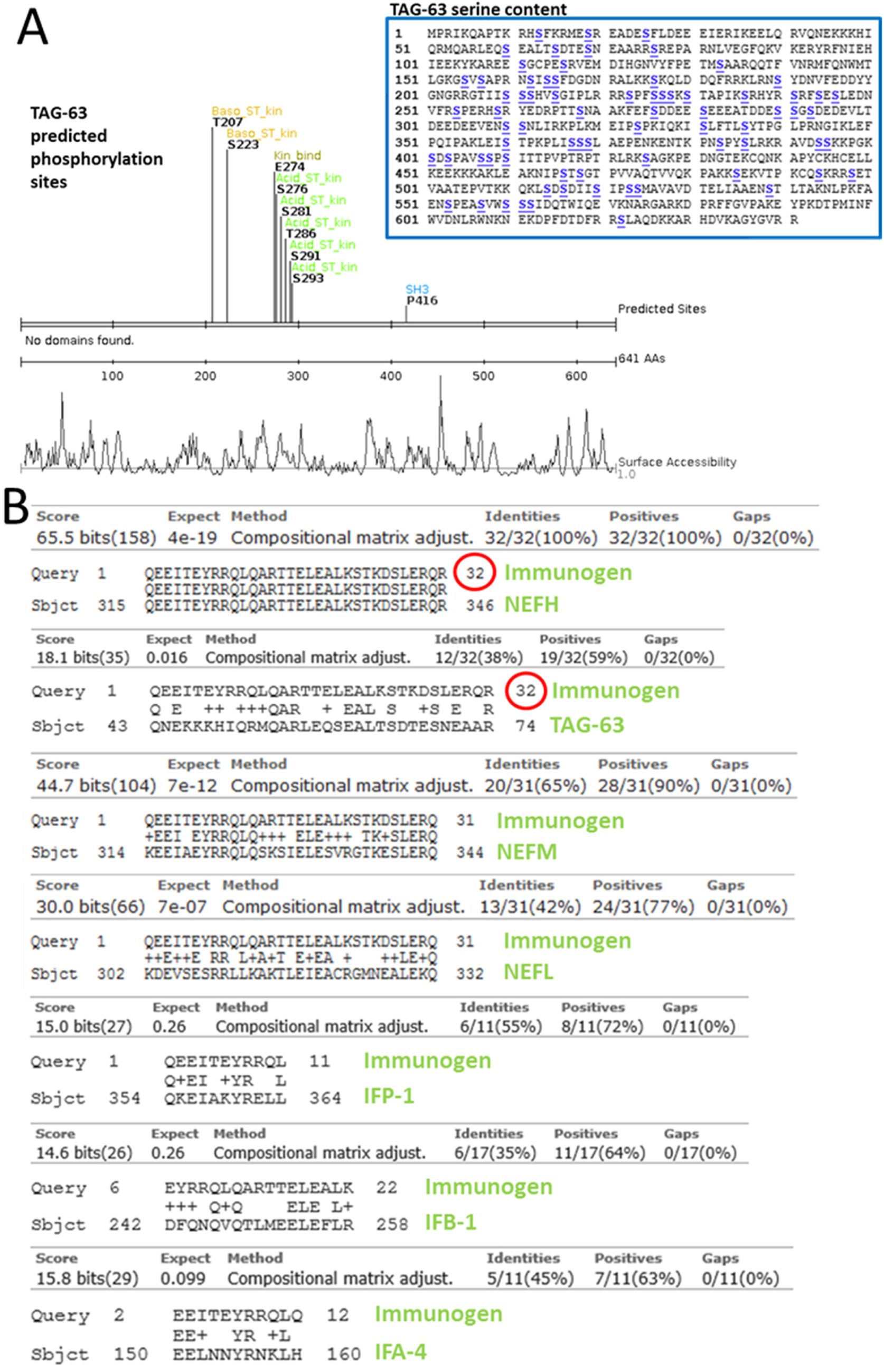
Prediction of phosphorylation sites in TAG-63 and anti-NEFH epitope sequence BLAST. (A) SCANSITE3 bioinformatics tool predicts nine phosphorylation sites clustered between coiled coils (Suppl. Figure 1C) in TAG-63 (serine content noted in the box to the right). (B) The sequence of the epitope to the anti-NEFH antibody (#WH0004744M1, Sigma) used for Western blots was kindly revealed by the manufacturer allowing a BLAST against NEFH, TAG-63, NEFM, NEFL as well as various *C. elegans* intermediate filaments (IFs). From these results, we conclude that the antibody detects TAG-63 rather than other *C. elegans* intermediate filaments.

**Suppl. Figure S4:**
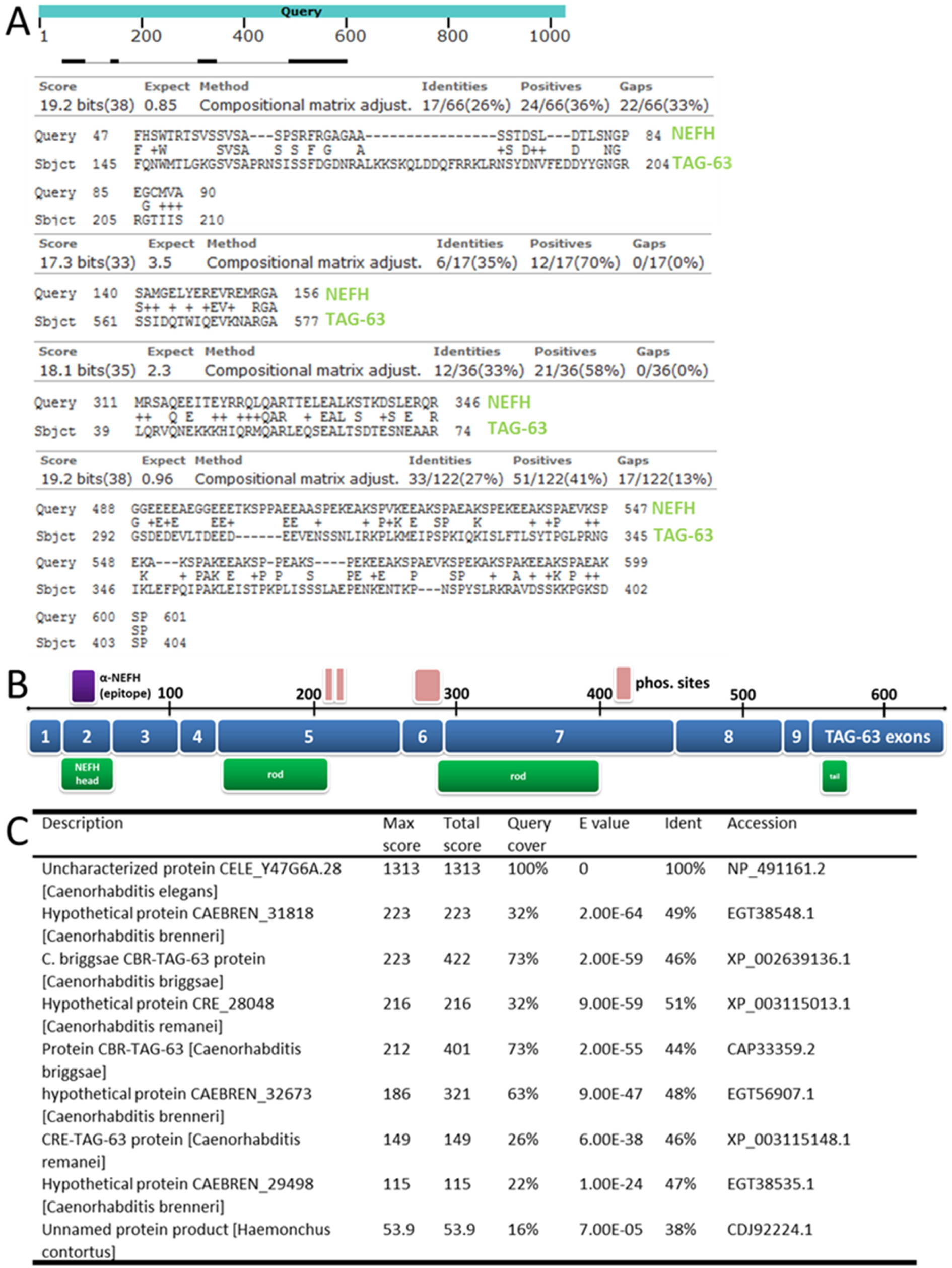
TAG-63 BLAST using NIH BLAST. (A) NIH BLAST using NEFH and TAG-63 as query reveals an average of 51% positives. (B) Schematic drawing of overlapping regions (blue is the TAG-63 sequence and green overlapping NEFH sequences). (C) Table of recognized genes using NIH BLAST with TAG-63 as input revealing various *C. elegans* TAG-63 paralogs.

**Suppl. Figure S5:**
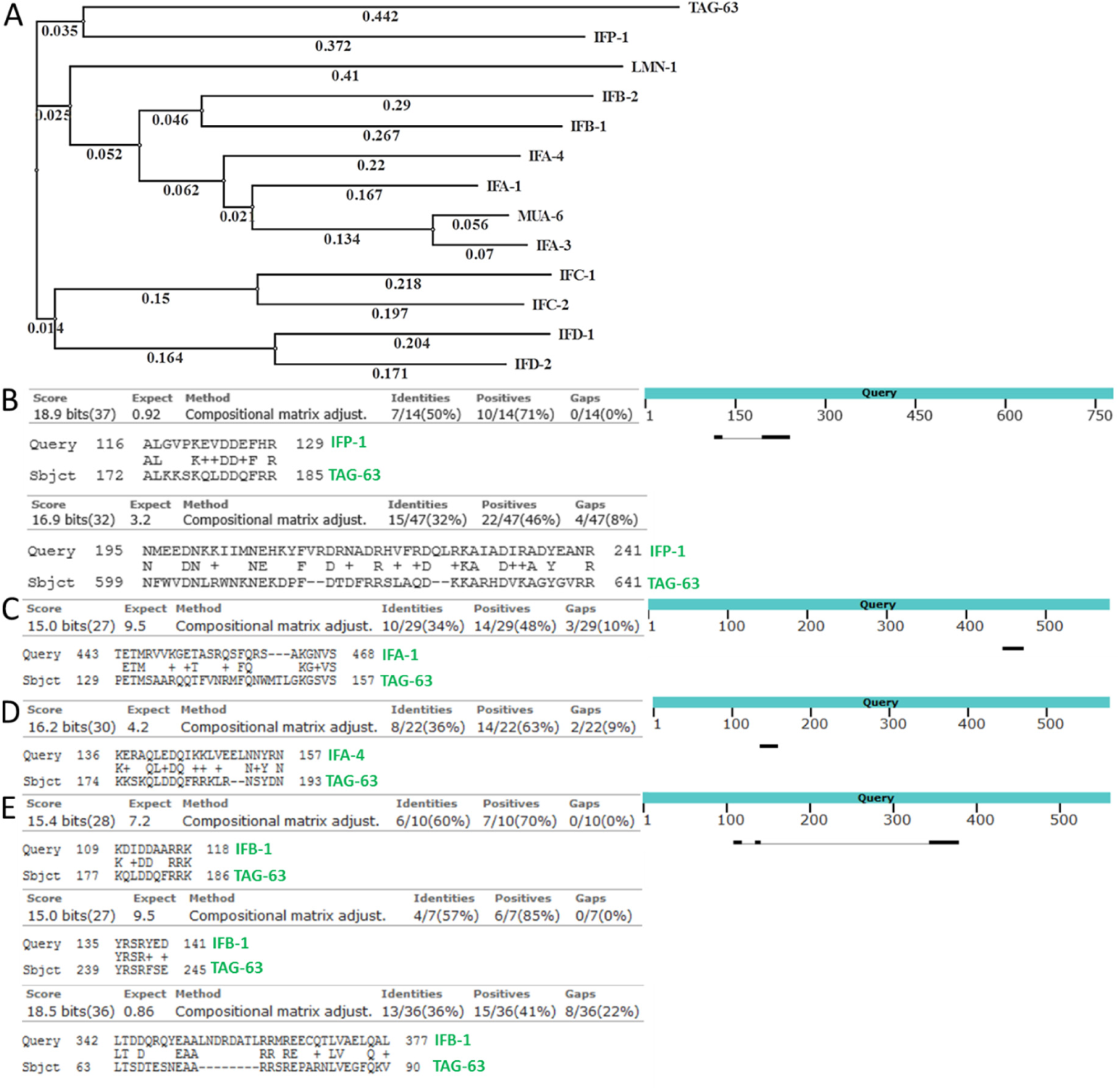
Phylogenetic tree and NIH BLAST analysis of TAG-63 against other *C. elegans* intermediate filaments. (A) Phylogenetic tree comprising TAG-63 and various *C. elegans* intermediate filaments reveals IFP-1 as being genetically close to TAG-63, however, sequence alignment (B-E) reveals only poor homologies to IFP-1 and other *C. elegans* intermediate filaments.

**Suppl. Figure S6:**
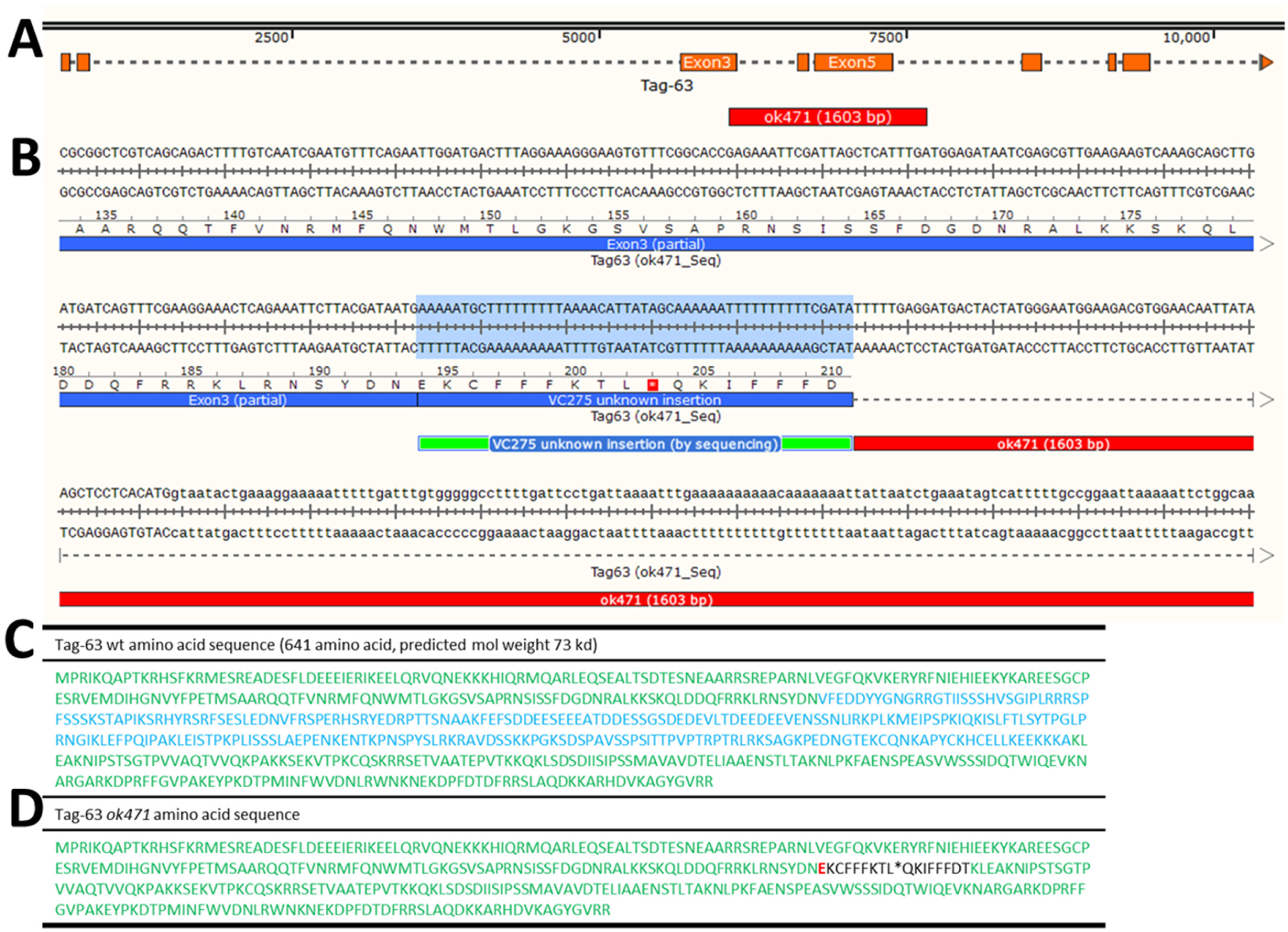
*Tag-63* genotyping. (A) The first eight exons and introns are shown in orange, and the annotated *ok471* deletion (from WormBase) marked in red. (B) Sequencing reveals (besides the annotated 1603 bp deletion) a 51 bp insertion at position 194 in exon 3 (green flanked region) with an amber stop (wildcard). (C) TAG-63 amino acid sequence with the *ok471* deletion area marked in blue. (D) Predicted amino acid sequence in *ok471* mutants with the deletion (annotated by WormBase) and the additionally identified insertion (marked in black) by us. The wildcard depicts the stop codon with the first amino acid at position 194 highlighted in red.

**Suppl. Figure S7:**
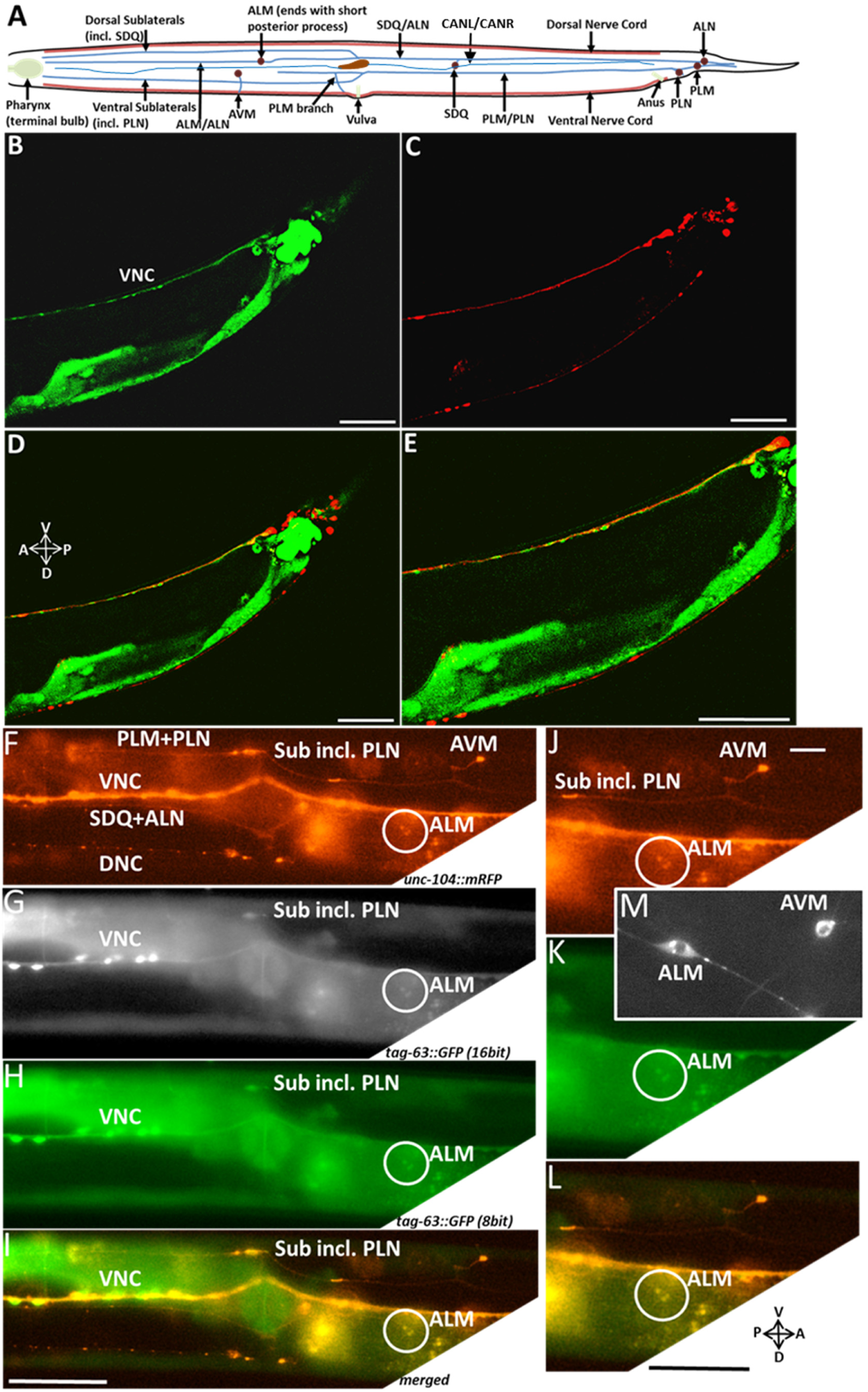
Expression pattern of *tag-63* transcriptional fusion overlapping with UNC-104 expression. (A) Schematic drawing of a simplified *C. elegans* nervous system. Anterior head (only terminal bulb of pharynx shown) is facing to the left and the posterior tail is facing to the right. Blue lines represent neurites and brown circles represent somas of neurons. (B) *Ptag-63::gfp* transcriptional fusion expression in the ventral nerve cord (VNC) near the tail of the worm. (C) Pan-neuronal *unc-104::mrfp* expression. (D) Merged image and (E) magnified image. (F-L) Overexposed images. In detail, (F) Pan-neuronal *unc-104::mrfp* expression (VNC, ventral nerve cord, DNC, dorsal nerve cord, sublateral neurons). (G) 16 bit grayscale and (H) 8 bit color image of P*tag-63::gfp* expression revealing neurons such as ventral nerve cord (VNC), sublateral neuron and ALM neuron and (I) merged image with *unc-104::mrfp*. (J-L) Magnifications from (F), (H) and (I). (M) ALM mechanosensory neuron (with its characteristic bipolar appearance and long posterior process) encircled in (J). Scale bars: (B-L) 50 µm, and (M) 5 µm.

**Suppl. Figure S8:**
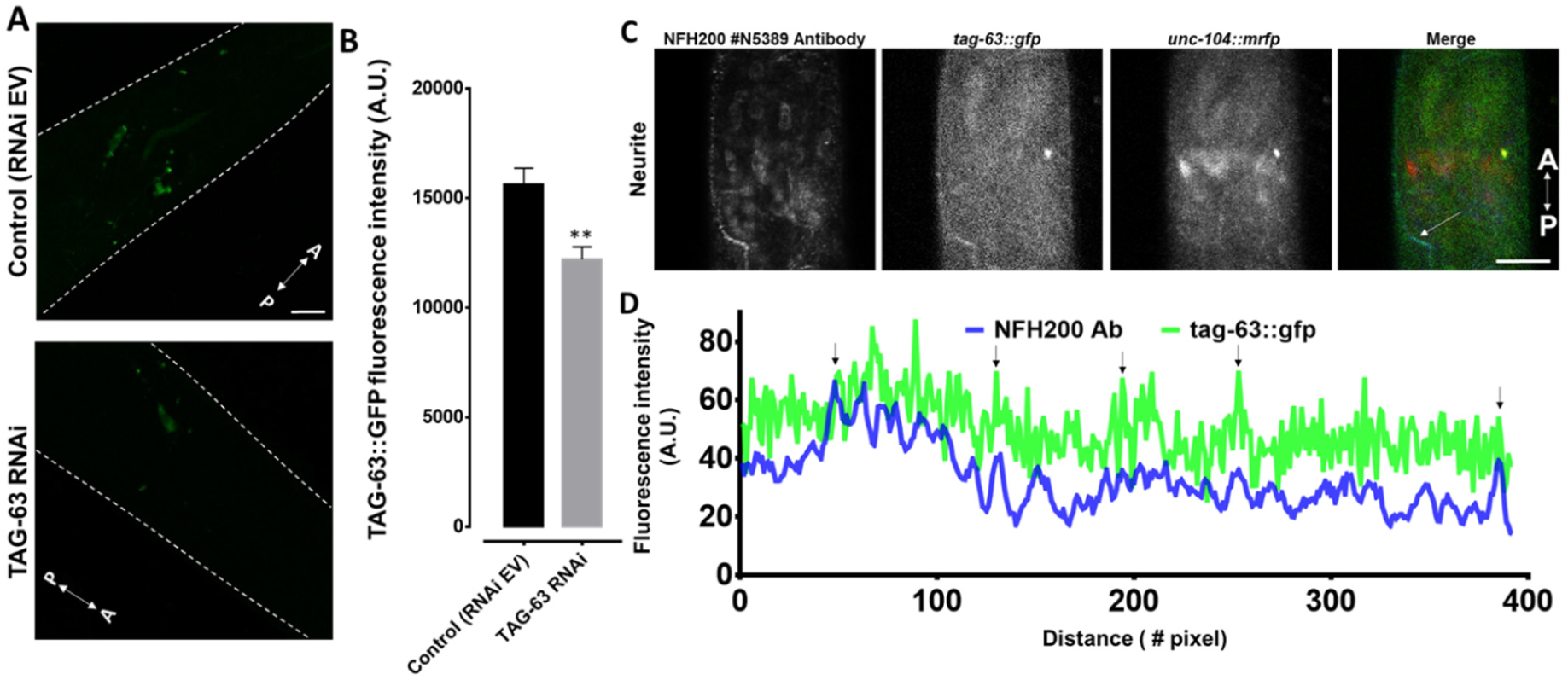
TAG-63 expression reduction after TAG-63 RNAi, anti-NEFH/TAG-63/UNC-104 triple staining. (A) Images of worm expressing *tag-63::gfp* treated with either empty vector (L4440, control RNAi EV) or TAG-63 RNAi clone. Dashed white lines outline the worm as seen under bright field. (B) Quantification of *tag-63::gfp* fluorescence intensity before and after TAG-63 RNAi. (C) Images of a worm immunostained with NFH 200 (#N5389) antibody co-expressing *Ptag-63::tag-63::gfp* and *Punc-104::unc-104::mrfp*. Merged panel: Blue: NFH staining, green: *tag-63::gfp*, red: *unc-104::mrfp*. White arrow indicates a representative neurite. (D) Line scan from the representative neurite shown in (C). Arrows point to overlapping peaks. Scale bars: (A+C) 10 µm. A: anterior direction and P: posterior direction. A.U.: arbitrary unit. **P<0.01 (T-test with Welch’s correction). Error bars: ± SEM.

**Suppl. Figure S9:**
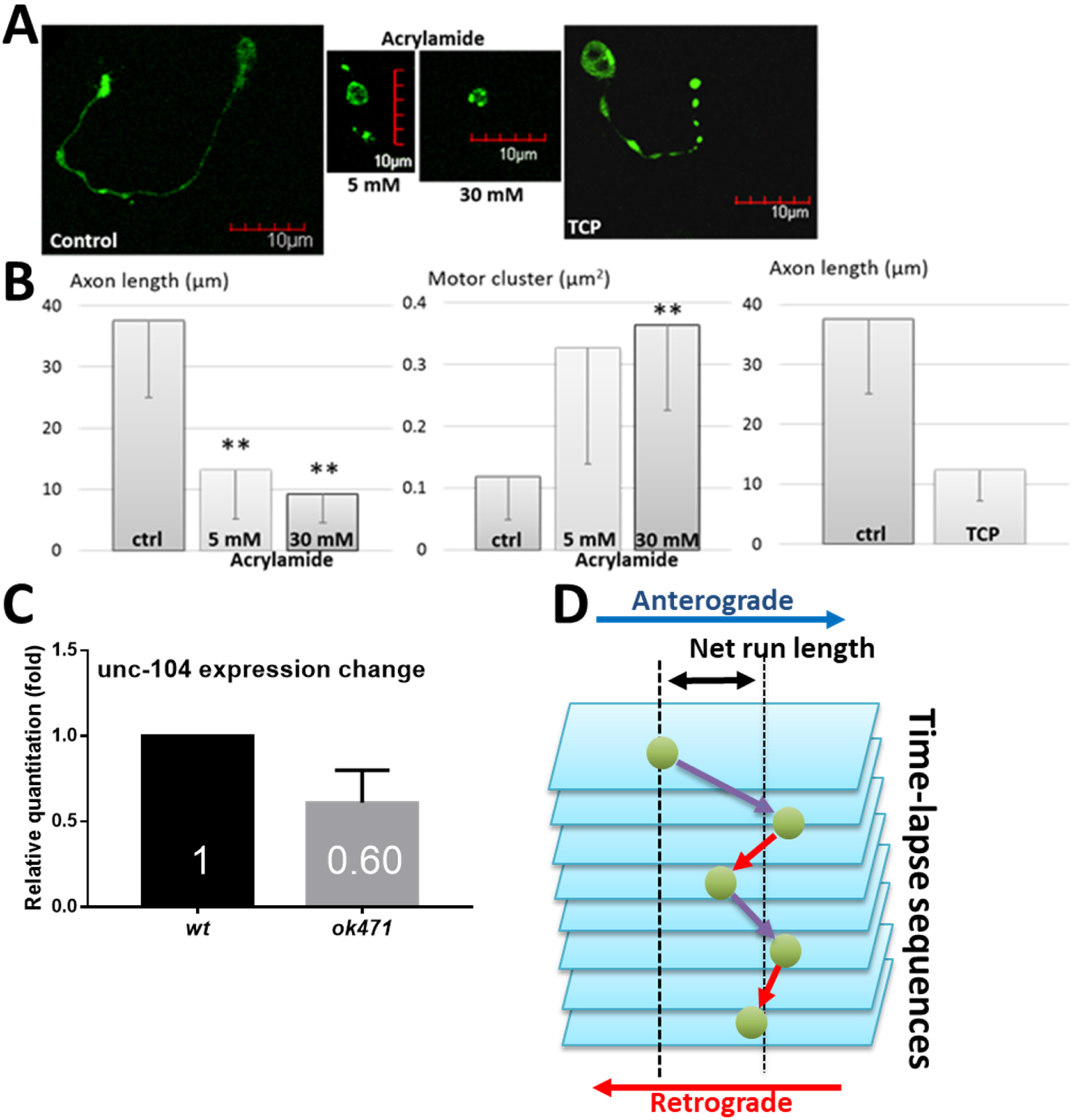
Treatment of cultured neurons with acrylamide and TCP (tricresyl phosphate), mRNA expression level and definition of used terms for motility analysis. (A) Treatment of cultured neurons (isolated from worms expressing *unc-104::gfp*) with different concentrations of acrylamide or TCP. (B) Quantification of axon lengths (and UNC-104 accumulations (cluster size)) in either acrylamide or TCP treated neurons. (C) UNC-104 mRNA expression level in wild type and *tag-63(ok471)* mutant worm. (D) Single anterograde moving events are represented by purple arrows, and single retrograde moving events are represented by red arrows. Flux is described as the number of events per minute. The double-headed arrow indicates the net run length of the green particle. Total anterograde run length is the sum of all purple arrows, and the total retrograde run length is the sum of all red arrows. Single event run length is distance covered by the particle in a single event. Scale bars: 10 µm. Error bars: ± SEM. **P<0.01(T-test with Welch’s correction).

**Suppl. Figure S10:**
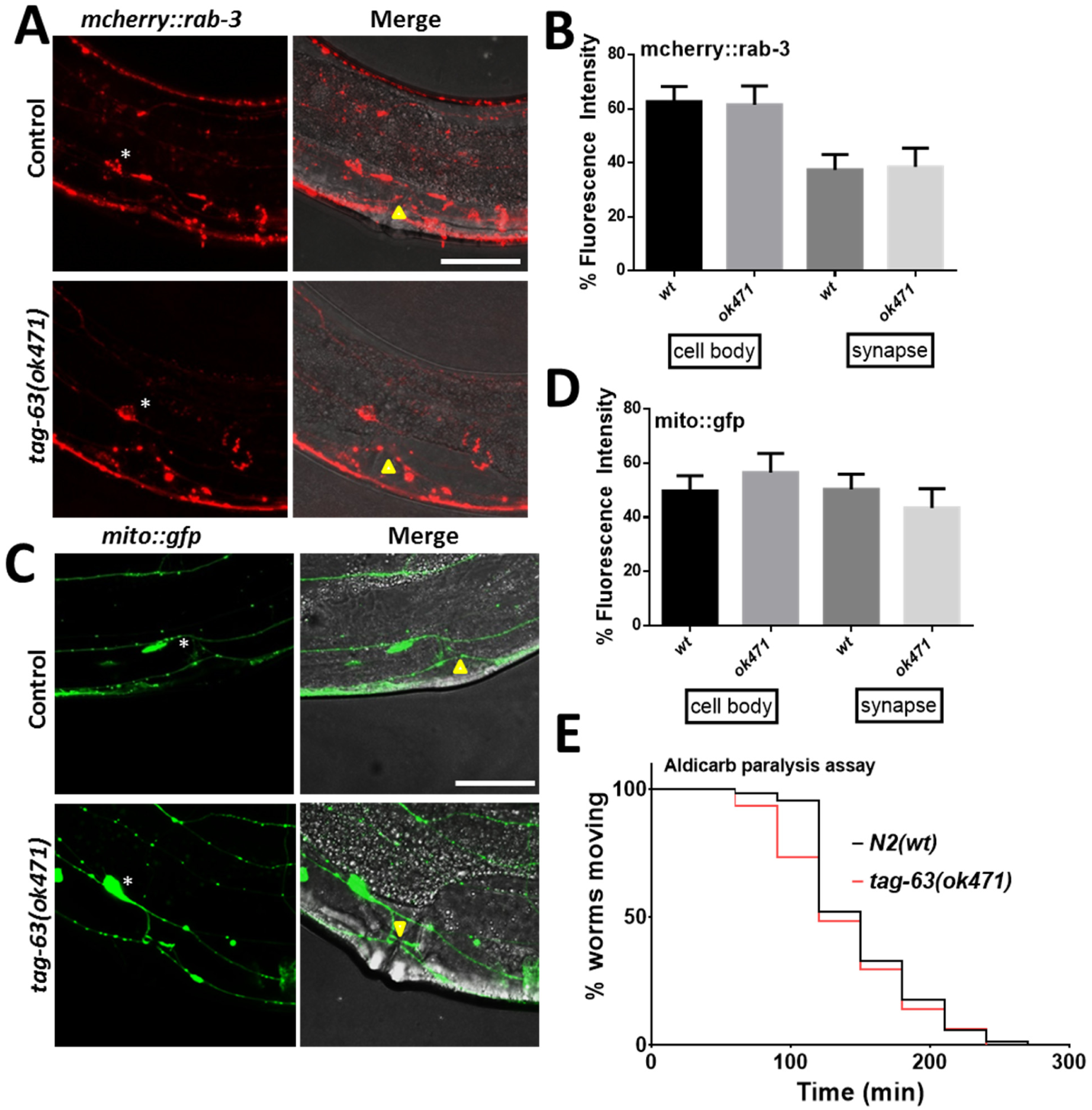
RAB-3 and Mitochondria transport in HSN neuron. (A) *Mcherry::rab-3* expression in soma and synapses of HSN neurons (wild type and mutant). (B) Quantification of *mcherry::rab-3* fluorescence intensity in the somas and synapses of HSN neurons as shown in (A). (C) *mito::gfp* expression in soma and synapses of HSN neurons (wild type and mutant). (D) Quantification of *mito::gfp* fluorescence intensity in the somas and synapses of HSN neurons as shown in (C). Yellow arrowheads point to vulva and wildcards “*” indicate the location of somas. (E) Aldicarb paralysis assay in wild types and mutants. Scale bars: 10 μm. Error bars: ± SEM.

**Suppl. Figure S11:**
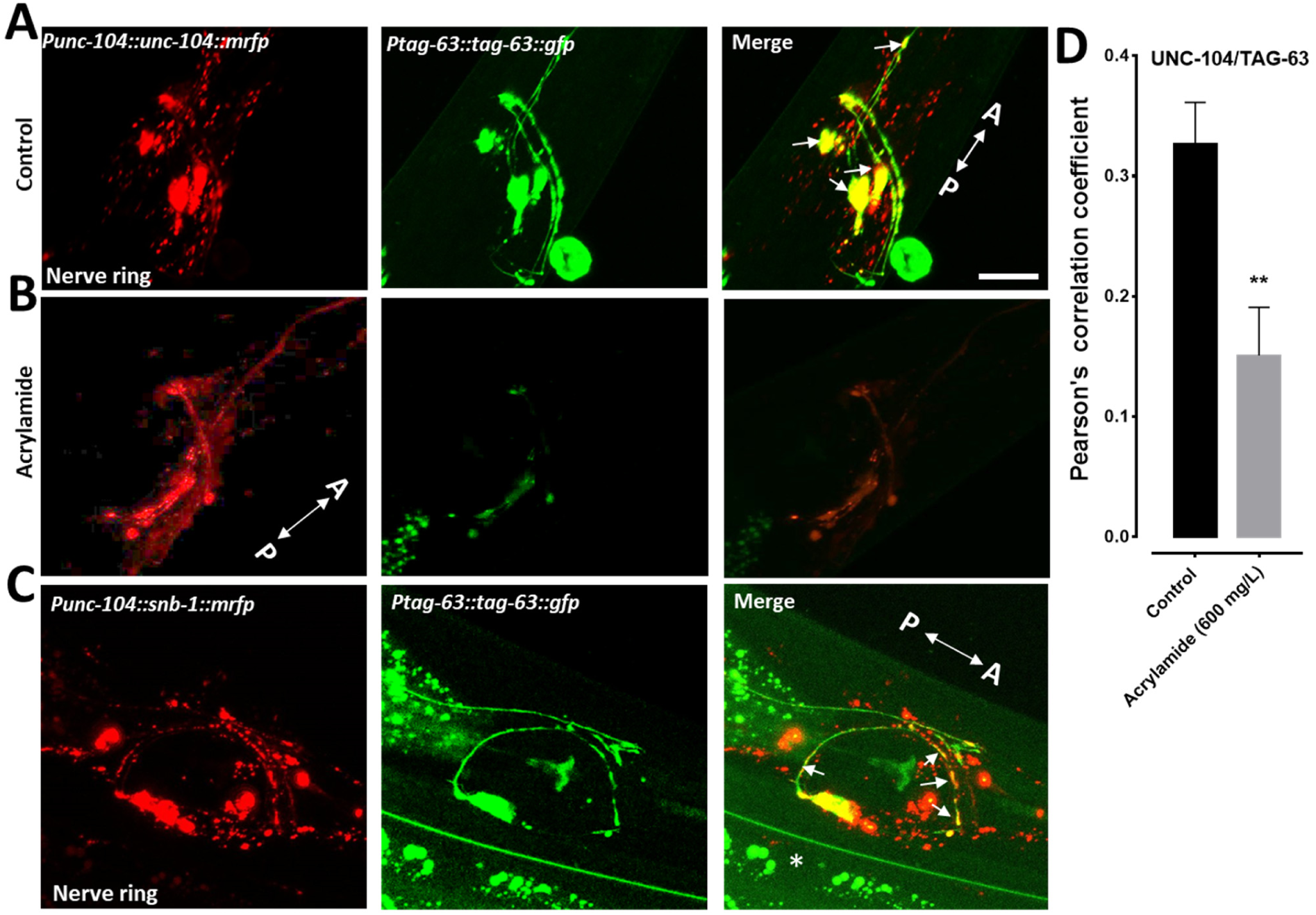
Colocalization of UNC-104 and SNB-1 with TAG-63. (A) Representative images of *unc-104::mrfp* colocalizing with *tag-63::gfp* in the nerve ring. (B) Worms co-expressing *unc-104::mrfp* and *tag-63::gfp* treated with acrylamide. (C) Images of *snb-1::mrfp* colocalization with *tag-63::gfp* in nerve rings. (D) Quantification of UNC-104/TAG-63 colocalization using Pearson’s correlation coefficient from (A+B) in control and acrylamide treated worms. A: anterior direction, P: posterior direction. White arrows (A+C) indicate colocalization signals. Error bars: ± SEM. Scale bar: 10 μm. **P<0.01 (T-test with Welch’s correction).

**Suppl. Video S1:**
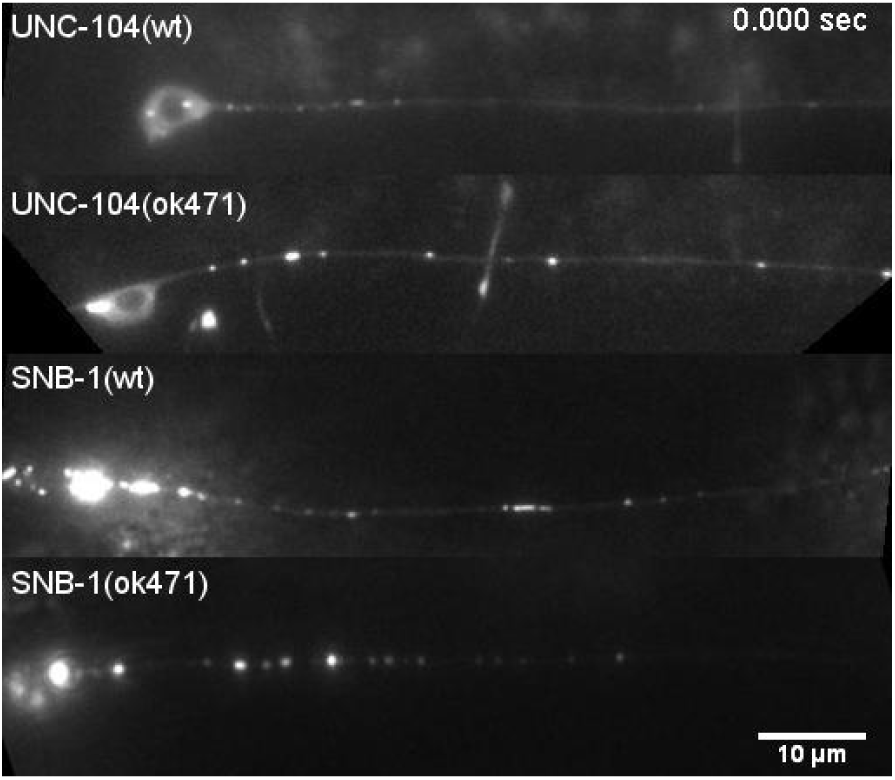
Representative videos taken from ALM neurites of worms expressing either *unc-104::mrfp* or *snb-1::mrfp* (*wt* compared to *tag-63* mutants). Increased clustering of UNC-104 in *ok471* mutants is obvious, while increased retrograde velocities, as well as reduced travel distances, are apparent for SNB-1 in *ok471* mutants. Video frame rate: 30 fps.

## Referee: 1

*We appreciate this reviewer’s valuable insights that greatly aided in improving the quality of our submission.*

### Comments to the Author

Bhan described the function of TAG-63 in axonal transport of *C. elegans*. Their results provide some valuable insights to the field. However, the following issues need to be resolved before publication anywhere.

1. The study relies on one single allele. There are numerous cases in which incorrect conclusions were drawn based upon one allele because of many reasons such as background mutation that is tightly linked to the locus. Considering the ease of CRISPR-Cas9, multiple alleles should be used to reach their conclusion. In addition, rescue experiments are essential.

*We thank this reviewer for pointing out this issue. Indeed, only one allele ok471 is available for tag-63, and we wish to emphasize that we did outcross the strain three times (see gels below) before employing it for any experiments. Additionally, we performed tag-63 RNAi experiments and resulting data are comparable to that of tag-63 knockout worms (see Figures 4, 6 and S8). Note that for all RNAi experiments we also carried out empty vector analysis as a control. Most critically, we overexpressed TAG-63::GFP in tag-63(ok471) mutant allele that successfully rescued UNC-104 velocities as well as net run length to wild type levels in tag-63 mutants (Figures 6 and 7).*

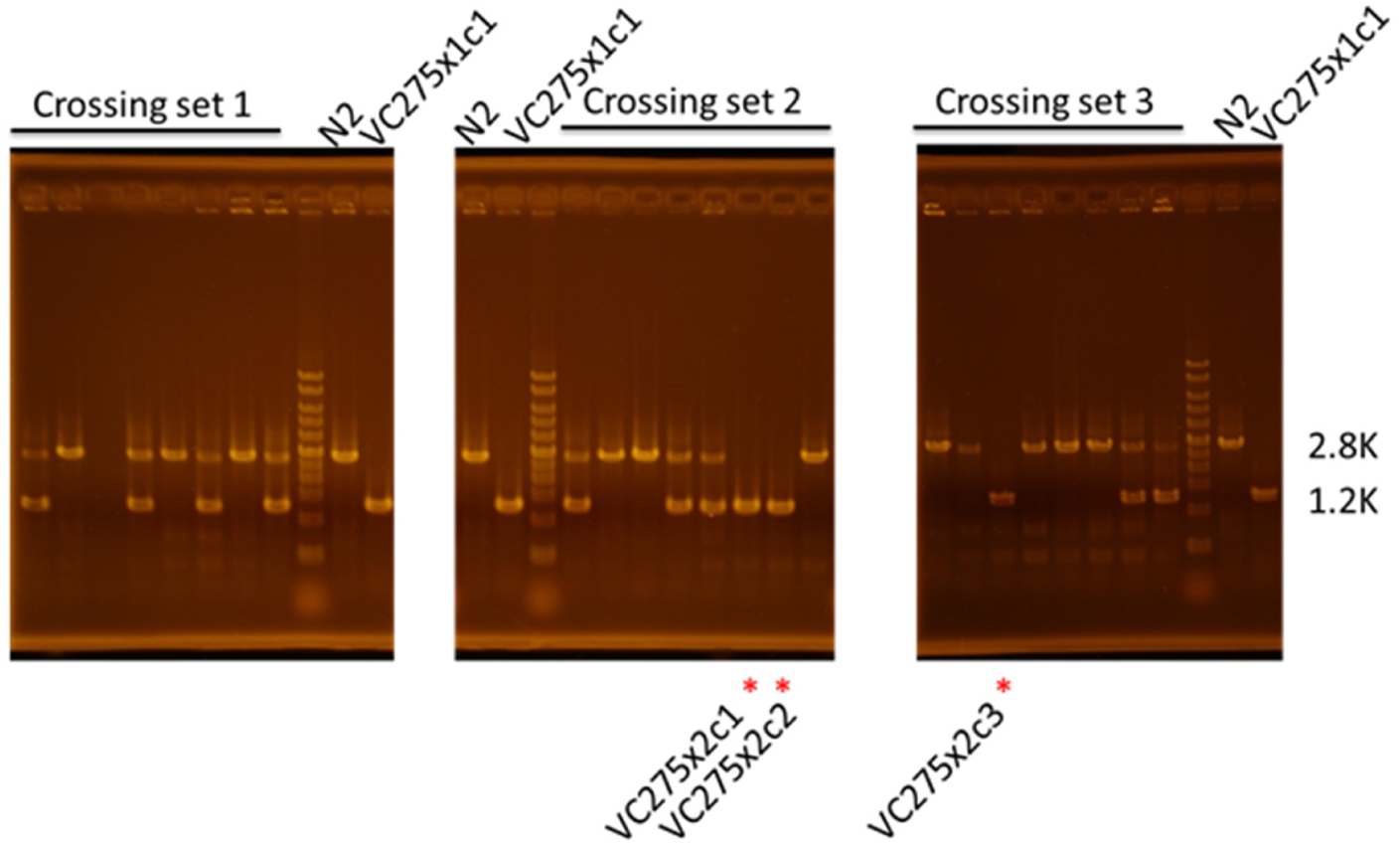

2. Many results are derived from a commercial antibody against mouse protein. While Western Blot in tag-63 deletion was performed, the identification of the identity of the proteins that were pulled down by this antibody should be subjected to mass spectrometry analysis. It is critical to validate the tool.

*As suggested by the reviewer, we have subjected the pull down proteins to mass spectrometric analysis and detected multiple binding partners (see table below). However, we did not detect TAG-63 even after three independent trials and it is not uncommon that a specific protein may be detected by Western blots but not be detected by shotgun proteomics such as LC MS/MS, specifically for larger and more complex proteins* ^*1*^. *Although Mass Spectrometry is extremely sensitive, protein signals might not be amplifiable to such extents as known for Westerns. Lastly, unlike purified TAG-63, in our experimental setup, we co-precipitated proteins (using NFH antibodies) from whole worm lysates and analyzed the bands around 73 kD, which is still a mixture of detected TAG-63 as well as of various other proteins residing in these bands.*

Requested Mass Spec experiment:

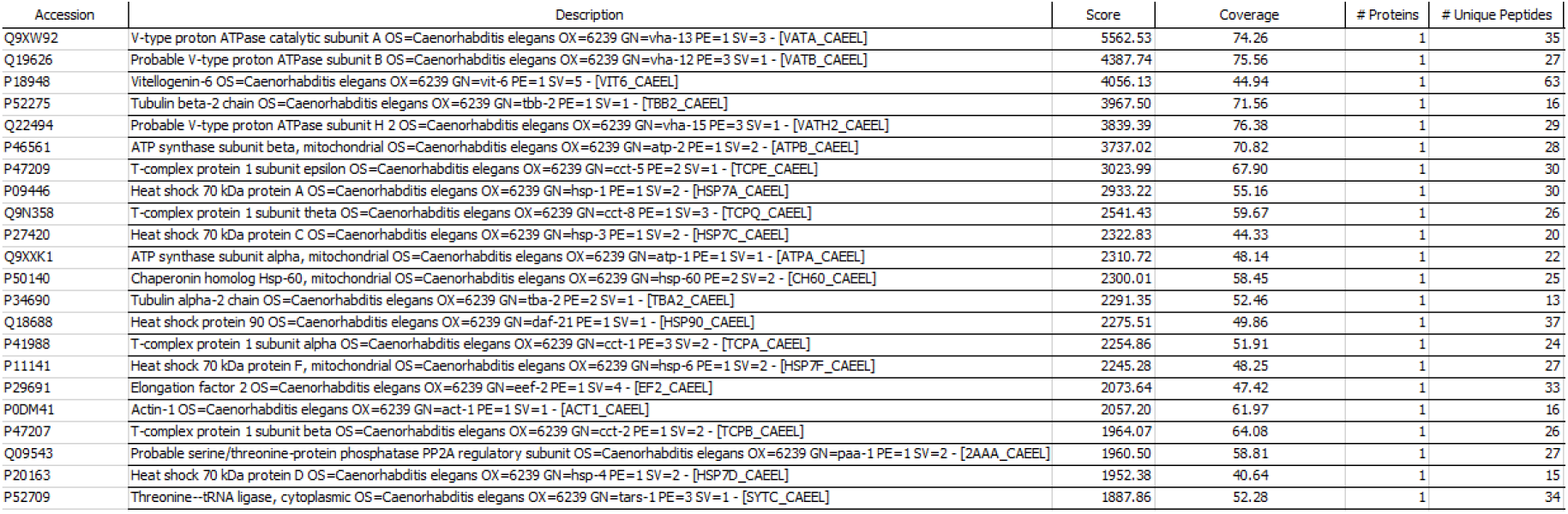

*Nevertheless, in order to further confirm the validity of our Western assays (as shown in Figure 1E+F), we now conducted another Western blot analysis using anti-NEFH, anti-NFH200 as well as anti-6X His tag antibodies to detect the recombinantly purified TAG-63 protein. All antibodies detected TAG-63 and all identified bands were located close to 73 kD matching the calculated molecular weight of TAG-63 (Figure 5B). As a reminder, please note that we provide BLAST sequence homology comparisons between the anti-NEFH epitope (#WH0004744M1, Sigma) and TAG-63 as well as to various other IFs in C. elegans* (Figure S3B). *These data firmly validate that the anti-NEFH antibody detects TAG-63 and very likely not other worm IFs.*

*New Figure 5B: Various NFH antibodies detect recombinantly expressed TAG-63 protein in Western blots.*

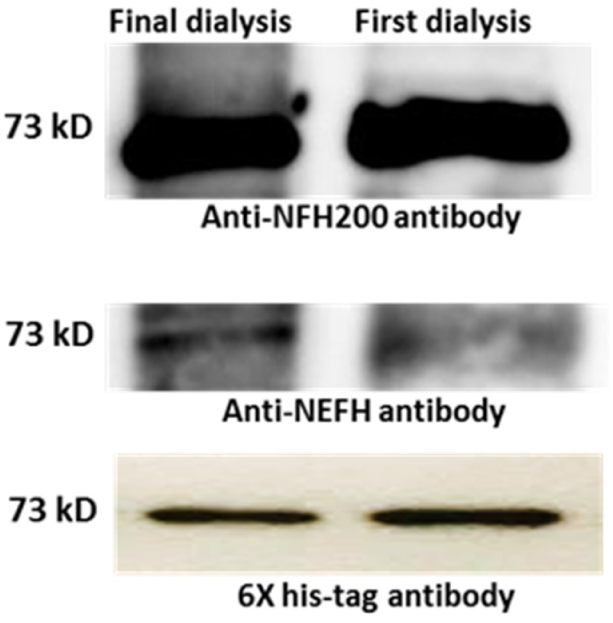

*Figure S3B: Sequence homology comparisons between a NEFH (#WH0004744M1, Sigma) antibody’s immunogen and C. elegans’ TAG-63 as well as other C. elegans’ IFs.*

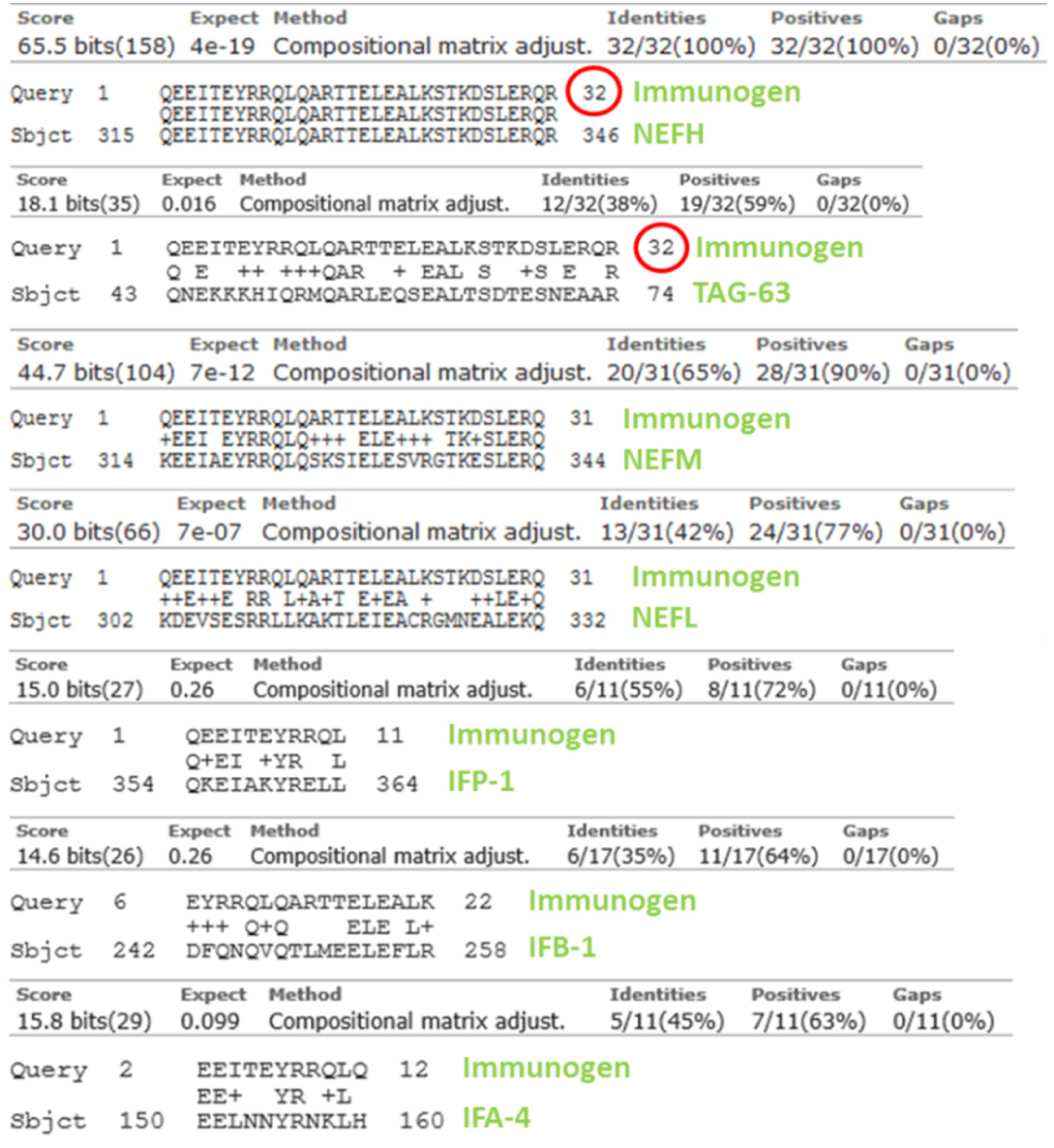

3. Figure 2 B * shows statistical significant. Is this really different?

*Based on our statistical analysis (student’s T-test), the data sets in Figure 2B indicate significant differences from each other with a p-value of 0.0473.*

4. Figure 4, the expression level change should be monitored using GFP knock-in.

*The reason why we analyzed mRNA expression levels of tag-63 gene in wild type and as well as in ok471 backgrounds (old Figure 4F, now new Figure 1D) is that we suspected a possibility of expression of truncated tag-63 gene in ok471 based on genotyping results as shown in Figure S6. Here we detect endogenous tag-63 gene and we did not use any strain expressing TAG-63::GFP.*

5. Figure 5, the EM experiments should be repeated. It is a big concern whether filaments rather than aggregates were formed.

*With great effort, we recombinantly expressed and purified TAG-63 and observed its filament-forming capacities using TEM revealing comparable structures as seen for neurofilaments by others* ^*2,3*^ *(see panel below). TAG-63 underwent expected structural changes upon urea solubilization and appeared as globular structures of sizes ranging from 120 to 180 nm. These globular units convert to either thick aggregates or fibrous bundles/filaments upon dialyzing against 20 mM Tris-buffer as shown by others for NFs. We performed experiments twice and results are comparable to the structures as described previously:*

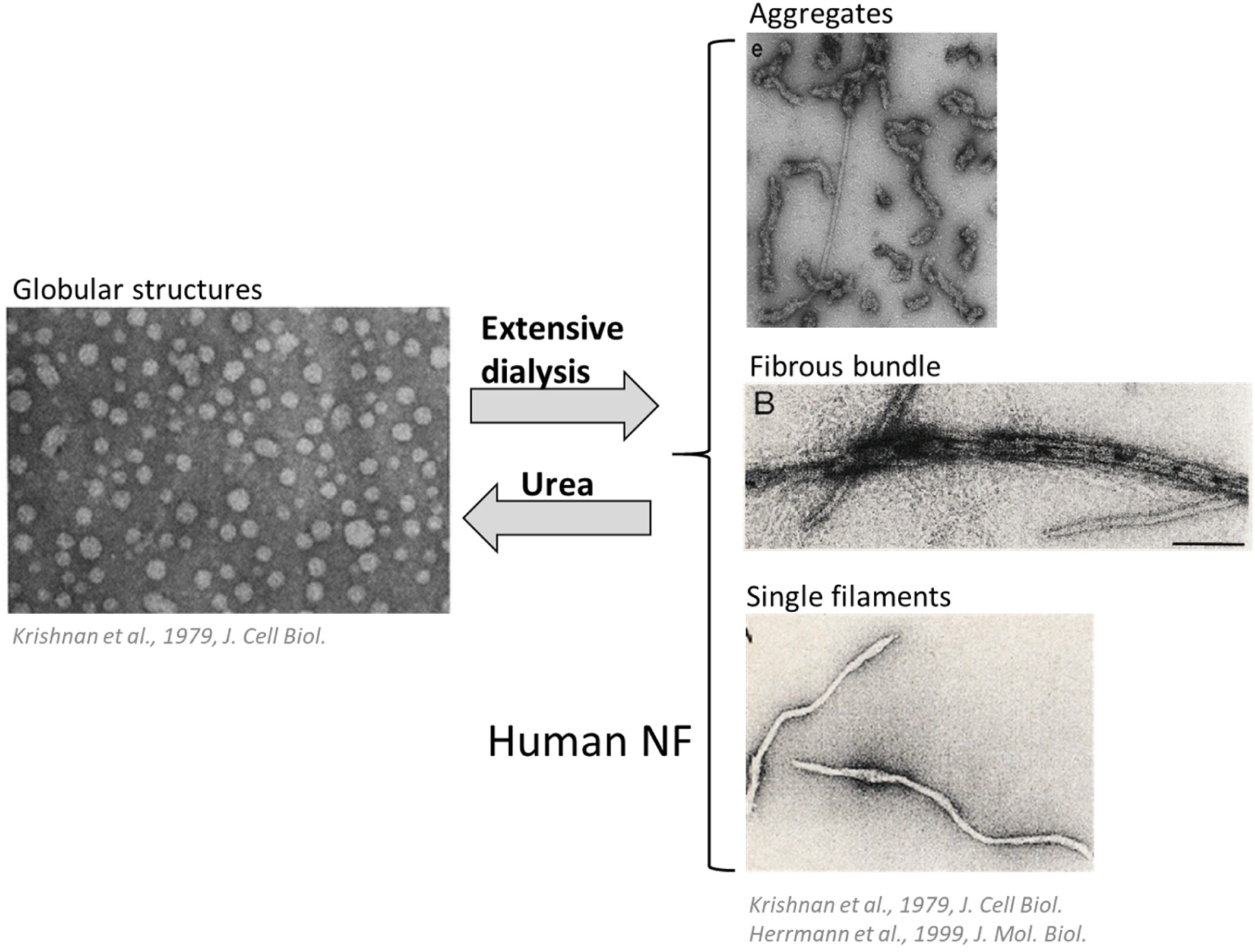

*Critically, we also visualized rhodamine-labeled (purified) TAG-63 appearing as filamentous structures with morphological similarities to rhodamine-labeled NFs observed by Wagner et al. 2003* ^*4*^. *These TAG-63 filaments easily disintegrate after exposing to the neurotoxin acrylamide (known to disintegrate NFs) in a concentration dependent manner (Figure 5G).*

6. The reduction of UNC-104 motility is really minor based upon speed and kymograph.

*We agree with the reviewer that UNC-104 anterograde velocity displays only one-star significant change in ok471 mutants, i.e., a minor reduction is observed. However, this reduction can be also seen after TAG-63 RNAi in wild types underlining the robust effect of TAG-63 on UNC-104 motility. More important, not only motor, but also vesicle trafficking is largely affected, as seen for SNB-1 travel distances and run lengths in Figures 7 and 8.*

7 It can be certainly helpful if TEM can be performed to visualize the neuronal phenotype at the ultrastructure level.

*We understand the curiosity of the reviewer in analyzing changes in sub-neuronal structures. However, our primary goals were to a) characterize TAG-63 as a novel NF orthologue in C. elegans and b) its role on axonal transport in C. elegans neurons. Future studies certainly may include the dissection of neuronal phenotypes by TEM; however, currently, such tedious and time-consuming additional experiments are certainly beyond the scope of our study.*

## Referee 2

*We thank the reviewer for thoroughly inspecting our submission and providing us important suggestions leading to an essentially improved manuscript.*

### Comments to the Author

#### Summary

Authors have characterized tag-63 and discovered homology to mammalian neurofilament-heavy polypeptides. Tag-63 is expressed in most C. elegans neurons and can form filaments. Loss-of-function of tag-63 shows increased accumulation of UNC-104 & SNB-1 in neuronal cell bodies and processes. TAG-63 also promotes anterograde transport of UNC-104 and UNC-104dependent cargo (SNB-1). Authors also claim that TAG-63 colocalizes with SNB-1 and UNC-104 in C. elegans neurons. They propose a model where TAG-63 supports the cargo-motor complex and promotes anterograde transport to synapses.

#### The main claims have not been substantiated

1) Authors claim TAG-63 colocalizes with UNC-104 & SNB-1: Images shown for colocalization analysis are taken at lower magnification and hence do not have sufficient resolution to claim juxtaposition of multiple markers. Additionally, UNC-104 shows a diffuse distribution-cannot do colocalization analysis between a punctate and diffuse marker.

*We thank the reviewer for carefully inspecting our Figures though we wish to point out that in high resolution images such as in Figures 8, 9E and VideoS1 UNC-104 reveals well-defined expression patterns in neurons. Also in kymographs (Figure 6 motility analysis) very well-defined lines can be seen that emerge from well-defined single UNC-104 punctae. We would also like to refer the reviewer to our previous publications*^*5,6*^ *for representative images on well-defined, non-diffusive and punctate UNC-104::mRFP/GFP expression patterns very similar as shown in this study (Figures 8, 9E and Video S1). The situation may indeed differ in nerve bundles or in nerve rings with overlapping somas and axons often leading to a clustered appearance of UNC-104. Also note that colocalization analysis was always performed within a selected region of interest to eliminate the possibility of insufficient resolution (see also Method section for further details on image analysis techniques).To further strengthen our colocalization data – and to comply with this reviewer’s concern - we now add new data from bimolecular fluorescence complementation (BiFC) assays (UNC-104/TAG-63 BiFC as well as SNB-1/TAG-63 BiFC) in new Figure 9D. These new and additional experimental data provide strong evidence for functional interactions between UNC-104/TAG-63 and SNB-1/TAG-63.*

*New Figure 9D: BiFC assays for UNC-104/TAG-63 and SNB-1/TAG-63 protein pairs:*

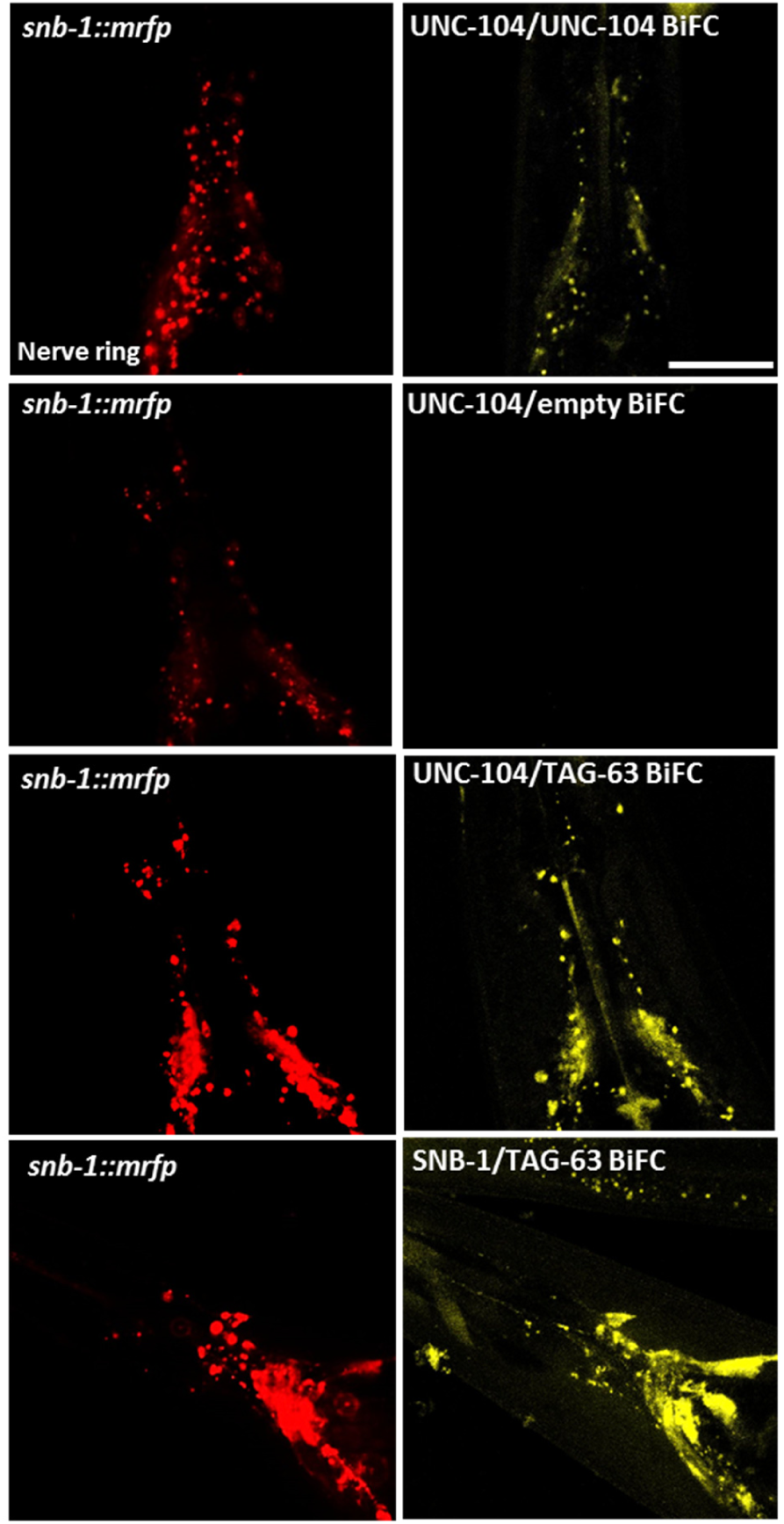

2) Authors claim TAG-63 promotes SNB-1 transport in an UNC-104 mediated manner: tag-63 mutants could have general morphological defects, thus causing accumulation of UNC-104 & SNB-1 along the neuronal process. Authors have not examined other motors and different cargo to examine whether these are general cargo defects or defects that affect specific cargo.

*We appreciate that the reviewer had raised this concern. We now add new data on accumulation studies for two other fast axonal transport cargos: RAB-3-containing vesicles and mitochondria. Interestingly, we observed that neither mCherry-tagged RAB-3 (known to be transported by kinesin-3/UNC-104) nor GFP-tagged mitochondria (known to be transported by kinesin-1) display any transport defects in HSN neurons; contrary to SNB-1 cargo that mislocalizes to cell bodies owing to compromised transport mediated by UNC-104.*

*New Figure S10A+B: RAB-3 transport is unaffected in tag-63 mutants*

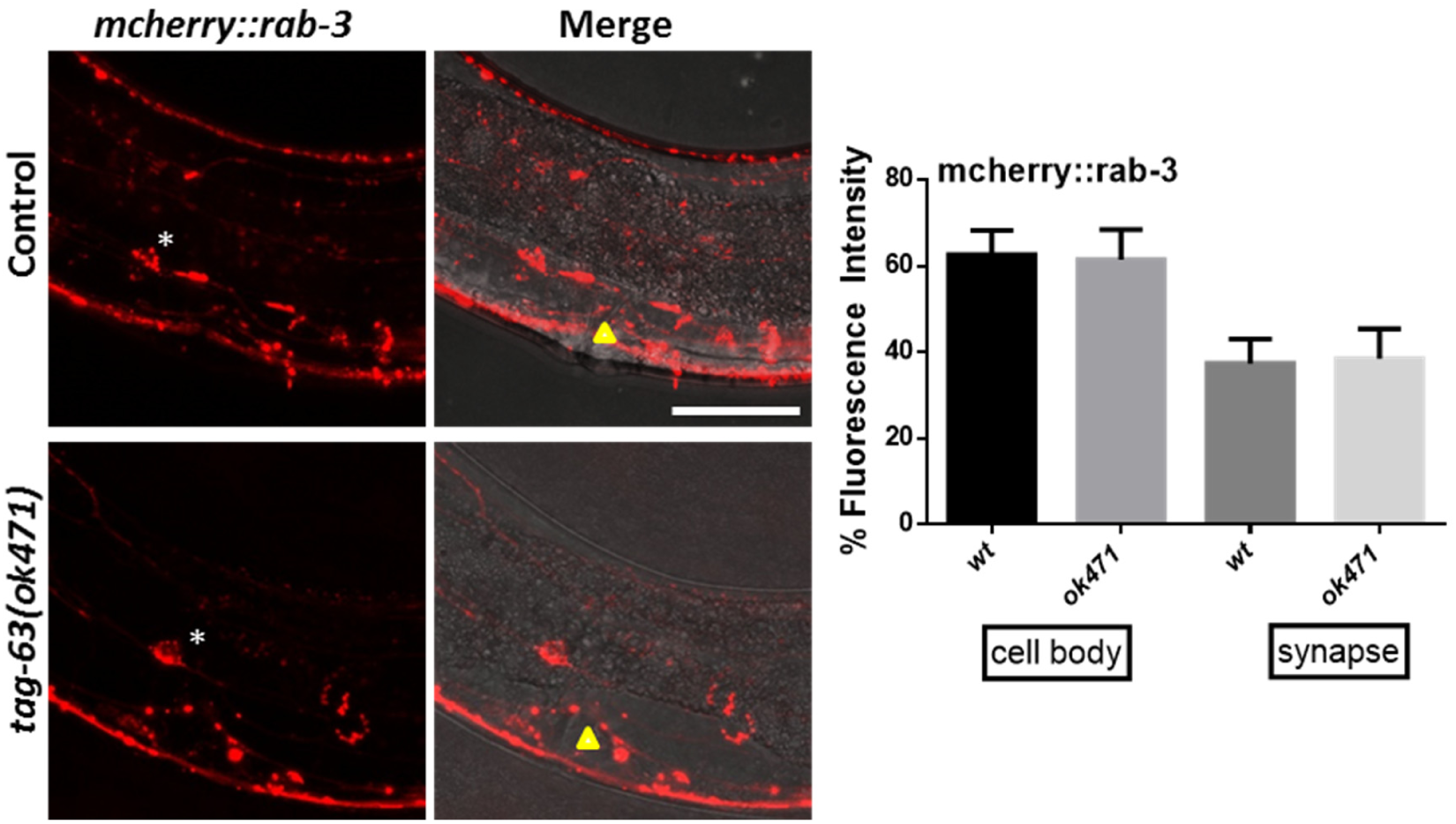

*New Figure S10C+D: No changes in mitochondria distribution in HSN neurons after tag-63 knock-out.*

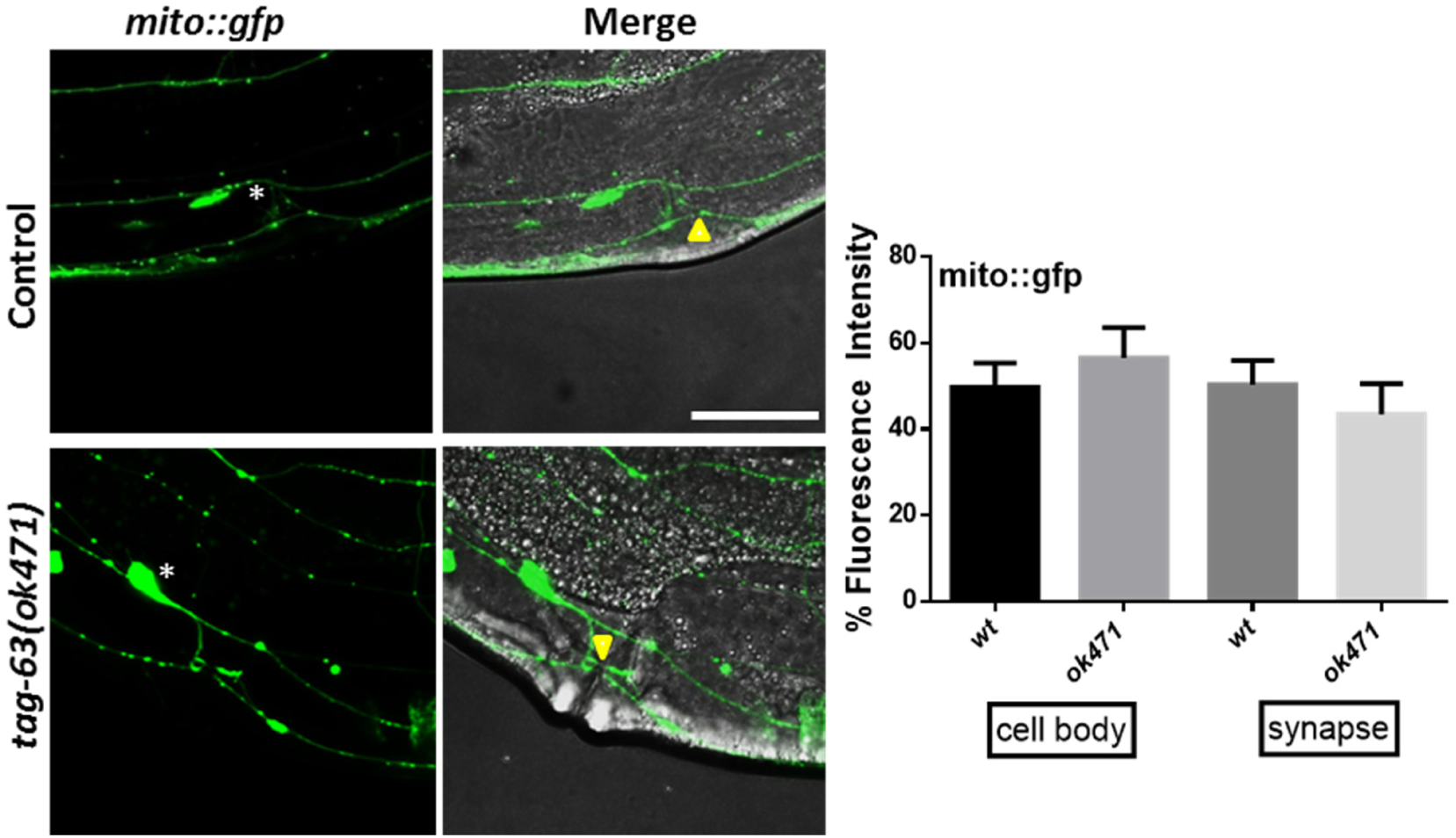

3) Authors claim vesicle transport to synapses is compromised in tag-63 mutants: Authors have not examined synaptic boutons to compare number of vesicles between wild type and mutants. Authors have not demonstrated any synaptic transmission defects in tag-63 mutants. Mutants only display mild egg retention phenotype and authors have not established a proper physiological significance to axonal transport defects seen in the mutant. One would expect wide spread cargo distribution defects and associated behavioral phenotypes.

*In C. elegans synaptic boutons are formed close to NMJs along the axonal shaft where the synapses often occur in an en passant manner between parallel nerve processes* ^*7*^. *As a new data addition, we now compare the number of synaptic vesicles between wild type and mutant at such synaptic boutons. As a result, we observed that in sublateral nerve cords*, SNB-1 *en passant puncta number was significantly reduced in tag-63 mutant worms as compared to wild type animals.*

*New Figure 8J*+*K: In sublateral nerve cords, the number of SNB-1 puncta is visibly reduced in tag-63 mutants.*

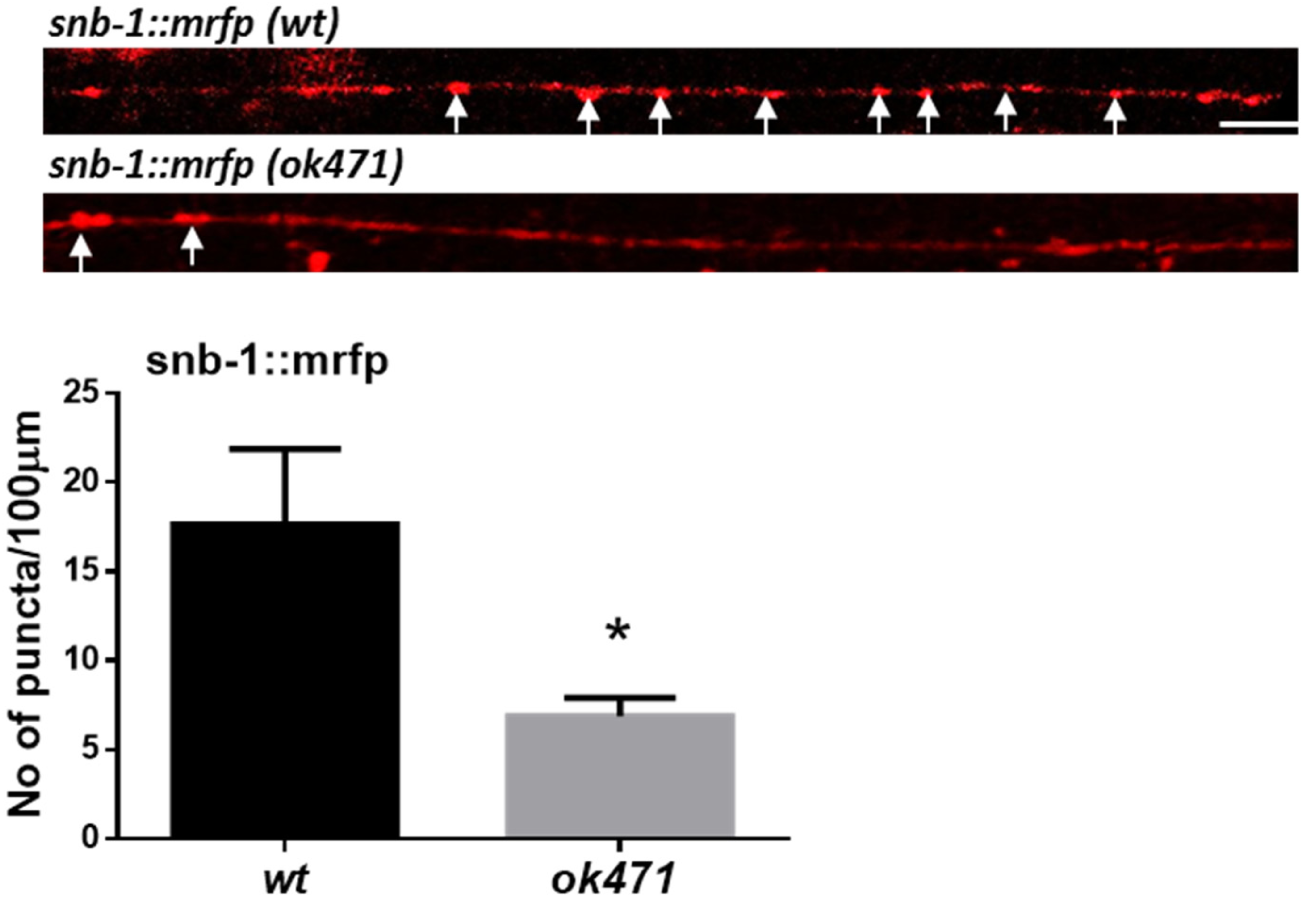

*Besides, to comply with the reviewer’s idea on investigating synaptic transmission defects, we have also performed aldicarb and levamisole paralysis assays (to understand whether or not synaptic transmission defects occur in tag-63 mutants). Animals with presynaptic defects, or lacking functional <u>acetylcholine receptors</u>, are known to be resistant to <u>aldicarb</u> while animals lacking functional <u>nicotinic acetylcholine receptors</u> display resistance to <u>levamisole</u>. Our new data provide evidence that tag-63 mutant worm’s displays sensitivity to aldicarb, however, this effect is tendentious and not significantly different from wild type animals. Although, SNB-1 transport is measurably affected in tag-63 mutants - as concluded from motility analysis as well as HSN neuron cargo accumulation analysis - we would indeed expect aldicarb resistance to a certain degree in tag-63 mutants. One explanation for the observed phenotype might be that normal transport of RAB-3 - as observed in tag-63 mutant worms (Figure S10A+B) - may compensate the SNB-1 transport defects to some degree. Strikingly, tag-63 mutants exhibited very obvious levamisole resistance, likely based on the downregulation of nicotinic acetylcholine receptors at NMJ’s as seen by others in various synaptic transmission mutants 8. Such strong synaptic transmission defects (Figure 2C) may be indeed related to defects in axonal transport of synaptic vesicles.*

*New Figure 2C+S10E: Aldicarb and levamisole paralysis assay*

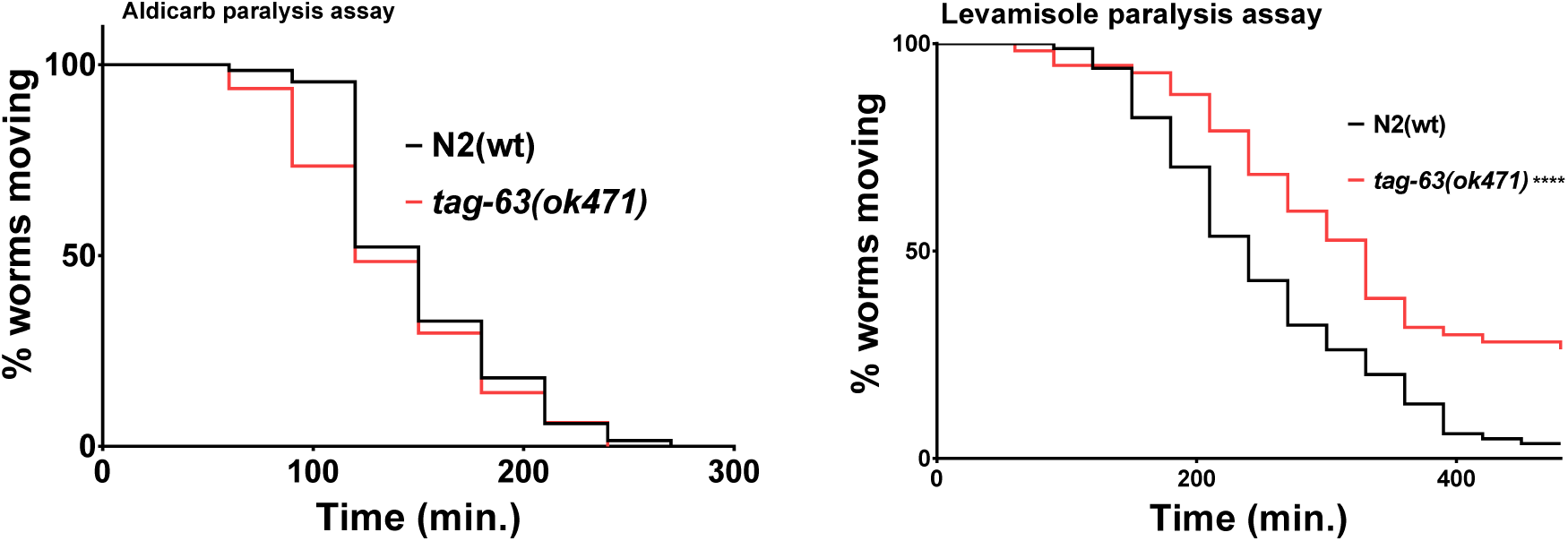

#### Technical comments

Genotype-phenotype analysis of tag-63(ok471) allele

Figure 2A: Representative images for wild type and mutant do not appear to be of the same stage. Are tag-63 animals larger in size than wt animals of the same stage? Bright field image of mutant-label should be corrected to tag-63(ok471). Figure legend mentions 10 worms were used for the egg-laying phenotype analysis, however, the graph shows over 20 data points for each genotype. Need more detailed description on what is plotted. T-test may not be the right test for this comparison as the data distribution does not appear to be Gaussian. A non-parametric test like Kruskal-Wallis may be better for comparison.

*The images of wild type and mutant belong to the same stage, however, due to egg retention phenotypes in tag-63(ok471) mutants, worms often appear bulkier. Also, as suggested by this reviewer the labeling has been corrected and non-parametric comparison test has been used for better comparison between wild type and tag-63(ok471) worms. Note that ∼10 worms were used for analysis in Figure 2B and in Figure 2A data points relate to the number of eggs retained for each analyzed genotype.*

Figure 2D: Since there appear to be only 3 animals used for the Chemotaxis analysis, Mean ± S.E.M. is the wrong way to represent this data for comparison. Additionally, the t-test may be inappropriate to use as three data points are too few to make a Gaussian fit. Kruskal-Wallis test may be better for comparison.

*We thank the reviewer for highlighting this issue. The three data points indicate chemotaxis indices from three independent trials each carried out with 19-25 worms. We revised the Figure legend accordingly.*

In addition to these characterization tests, it may be advisable to perform an aldicarb sensitivity assay to examine whether synaptic transmission is affected in tag-63 mutants. This may will also be a physiologically relevant assay considering the proposed model of tag-63 animals having fewer synaptic vesicles transported to synapses.

*Please refer to the above section in which we discussed the newly added data on aldicarb and levamisole paralysis assays.*

1) Neither the Figure 3 legend not the methods section mentions what objective was used for fluorescence imaging.

*A 100X objective lens on a Zeiss LSM 780 laser confocal microscopic system was employed for imaging worms. This information has been updated in the revised manuscript.*

2) For each panel, it is preferable to show a schematic of the neuron studied, to make it easier for general readers to understand what they are looking at.

*As suggested by the reviewer, we have now included a schematic representation of examined neurons in this study in new Figure S7A.*

3) The individual panels need to be better aligned and of the same size.

Figure 3C: No white arrows shown indicating common neurons expressing both UNC-104 and TAG-63

*Necessary changes have been made to the revised manuscript.*

Figure 4A & B: Co-localization of TAG-63 & UNC-104 is not very meaningful as UNC-104 distribution is diffuse throughout the cell. For co-localization analysis, images need to be taken at higher magnification (60X/100X). RFP and mCherry tags are known to clump in C. elegans. It is highly advisable to use UNC-104 GFP for artefact-free data comparison.

*Please note above our comments on “diffuse UNC-104 staining” and newly added data employing BiFC assays.*

*Besides, in Figure 4, for the whole mount immunostaining protocol worms must be subjected to reducing agents such as β-mercaptoethanol as well as collagenase incubation, both may affect normal protein distribution. Still, well-defined UNC-104::GFP punctae in Figure 4A as well as UNC-104::mRFP punctae in Figure 4B can be seen. Also please note that our opinion on the usage of red fluorophores in C. elegans slightly differs such as it has been clearly shown that monomeric forms of red fluorescence protein demonstrate high photostablity and GFP rather tends to dimerize and aggregates 9,10.*

Figure 4C & E: Fluorescence intensity calculations need to be internally normalized to a soluble fluorophore’s intensity, to compare across genotypes.

*The method to calculate the fluorescence intensity has been extensively described in the Methods & Materials section of the revised manuscript. The background fluorescence is subtracted from the integrated density of the region of interest and then the cell’s corrected total fluorescence is calculated using the formula*

*CTCF = Integrated density – (area of selected cell X mean fluorescence of background readings).*

*This method has been also employed in an earlier publication* ^*11*^ *to quantify immunostaining. Further, immunostaining detects the endogenously expressed protein TAG-63 in young adult worms or age synchronized worms using comparable dilution of antibodies across different genotypes. Further, we did not normalize fluorescence intensities for tested genotypes, because Figure 4C and 4E quantifications are not presented in the fold change unit.*

Further, using anti-tag63 Ab when expressing TAG-63::GFP, co-localization is obvious-not the right experiment to do.

*We used anti-NFH200 antibody to detect TAG-63::GFP and anti-TAG-63 antibody has never been used in this manuscript. The antibodies used have been explicitly stated in the Methods & Materials section of the revised manuscript.*

Effect of tag-63 on UNC-104 (motor) and SNB-1 (cargo) transport efficiency:

1) Authors have not shown experiments that rule out gross morphological defects in neurons, due to which non-UNC-104 cargo may also show same distribution defects.

*Most of our analysis has been conducted in ALM mechanosensory neurons. While conducting these experiments, we did not observe any visible morphological changes in these neurons (Figure 8A-D, stacked ALM neurons show no morphological defects). Further, we also did not notice any changes in HSN or sublateral neurons.*

2) Authors have not ruled out the possibility that UNC-104 levels may be reduced in tag-63 mutants, thus causing defects in SNB-1 localization.

*We appreciate this valuable comment. We add new data revealing no significant changes in UNC-104 mRNA transcripts in tag-63 mutants.*

*New Figure S9C: mRNA expression levels of UNC-104 in wild types and ok471 mutants.*

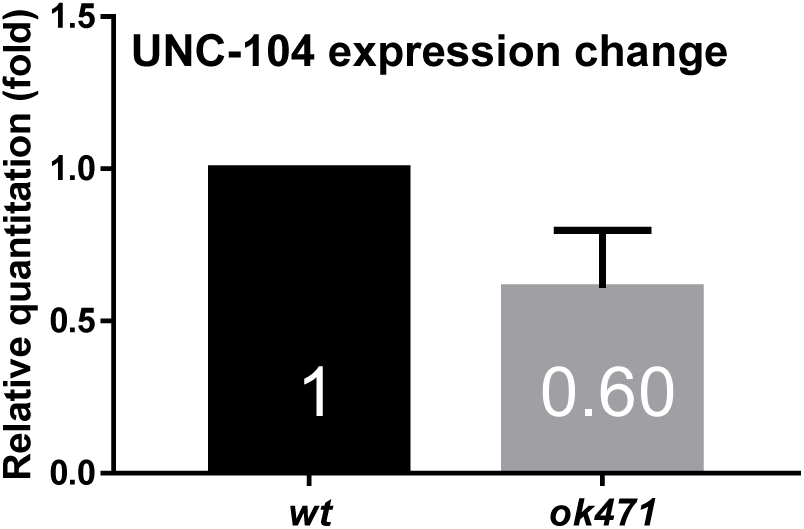

3) Authors state “it is apparent that UNC-104 motors move longer distances in a shorter period of time in mutants (as opposed to wild types) specifically obvious in retrograde directions” but in Figure 6A & B it is shown that UNC-104 anterograde velocity is significantly lower in the mutant as compared to wild type while retrograde velocity is comparable. In later paragraphs also, it is mentioned that UNC-104 is decelerated in the mutant, leading to increased accumulation.

*Moving distances are difficult to correlate to velocities. The reason why motors move longer or shorter distances relies on various parameters. First, molecular motors inherit their intrinsic duty cycles and dwelling phases. These may be shortened by obstacles on microtubules such as MAPs (tau etc.). Further, microtubules often have abrupt endings* ^*12*^ *(appearing half-staggered in C. elegans neurons) likely forcing the motor to detach and/or change MT tracks. Based on these issues we would refrain from correlating moving distances to velocities. Note that in a 2016 paper from our group* ^*6*^, *we investigated different moving speeds and distances in different regions of the neuron. For example, in the initial region of the ALM neuron UNC-104 motors move slower and shorter distances. While at more distal regions motors seemed to be more stably attached to MTs and running longer distances at higher speeds. In summary, moving distances as shown in the bullet diagram are the results of a complex interplay of various parameters (including duty cycle, dwelling phases, obstacles, MT distributions, tug-of-war events, active motor regulation by adaptor proteins etc.) all affecting the attachment- and detachment rates of motors.*

*Also, please note that UNC-104 is not a retrograde motor. In fact UNC-104 barely moves retrogradely in in-vitro motility assays 13. Thus, the retrograde component is reflected by the action of dynein and – though it needs further investigations in follow up studies – from our data it is obvious that in tag-63 mutants dynein-based retrograde SNB-1 transport is facilitated.*

4) In Figure 6: neither the legend nor the result section mentions which data has been acquired from primary cultured neurons and which has been acquired in vivo. If all the wt and mutant analysis is done by imaging the animals (and not primary culture), while acrylamide treatment is done on cultured neurons, then data from these two cannot be directly compared.

*We would like to bring to the reviewer’s notice that we have not acquired any data from primary cultured neurons in Figure 6. All wild type and mutant analysis has been conducted on whole animals. Negative affect of acrylamide on axonal transport has been reported by other groups* ^*14,15*^. *Note that worms can uptake acrylamide via NGM agar plates* ^*16*^. *Lastly, only data represented in Figure S9A+B have been acquired from neuronal cell cultures.*

5) In Figure 6A & B: appropriate control for acrylamide treatment on cultured neurons is missing.

*As mentioned above, we have not conducted our analysis in cultured neurons in Figure 6.*

In Figure 6C: UNC-104 kymographs look very similar SNB-1 kymographs and do not show any stationary UNC-104 particles. Do all UNC-104 moving puncta look like moving vesicles? If most of the moving UNC-104 particles are associated with vesicles, then shouldn’t SNB-1 show a reduction in anterograde velocity in the mutant?

*From Figure 6 at least one particle seems to be clearly stationary. Nevertheless, our analysis is based on (in average) 600 moving events per group and a single presented kymograph indeed often does not reflect the statistical nature and significance of the results. Further, from our newly added colocalization data (Figure 9E+F) it is evident that colocalization between UNC-104 and SNB-1 is significantly reduced in tag-63(ok471) mutant background likely due to the lack of functional support from TAG-63 filaments leading to variances in transport characteristics. More importantly, the reduction in anterograde run-length of SNB-1 is obvious from Figure 7G (although its anterograde velocity remained unchanged). Additionally, due to reasons mentioned above, the retrograde component of SNB-1 is likely reflected by the action of dynein and – though it needs further investigations in follow up studies – from our data it is obvious that in tag-63 mutants dynein-based retrograde SNB-1 transport is facilitated.*

*New Figure 9E+F: Colocalization of UNC-104/SNB-1 in sublateral nerve cords of wild type and tag-63 mutants.*

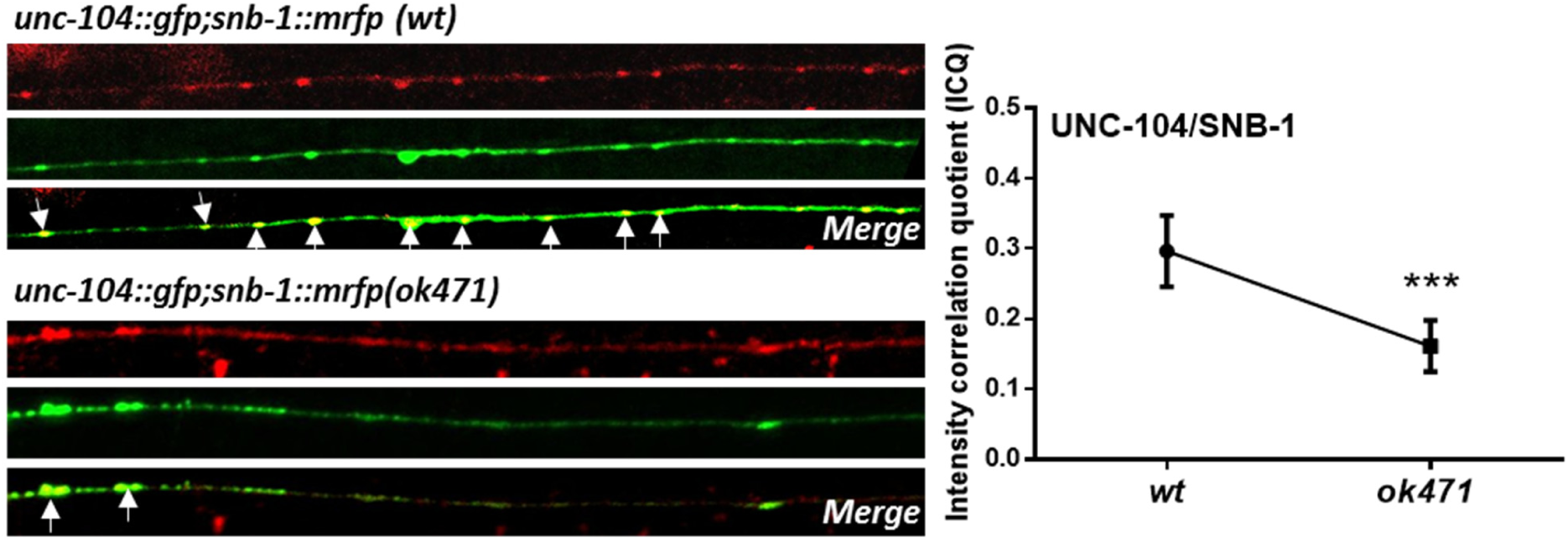

7) In Figure 6 & 7: UNC-104 transport characteristics do not map onto SNB-1 transport characteristics. It is unclear how the authors claim that tag-63 mutants show defects in axonal transport by affecting UNC-104 mediated transport.

*Kindly, refer to the explanation provided in reviewer’s point 3 and 6.*

Abnormal targeting of synaptic vesicle precursor (SNB-1) in tag-63(ok471) mutants:

Figure 8A,B,C & D: All the figures need to be better aligned and of the same length.

*We have modified the revised manuscript accordingly.*

Figure 8I: Intensity quantification in HSN neurons-no internal control used for normalizing fluorescence intensity, hence comparison across genotypes is not accurate.

*As mentioned above, the method to calculate the fluorescence intensity has been now extensively described in the Methods & Materials section of the revised manuscript. The background fluorescence is subtracted from the integrated density of the region of interest selected and then the cell corrected total fluorescence is calculated using the formula:*

*CTCF = Integrated density – (area of selected cell X mean fluorescence of background readings).*

*This method has been used in earlier publication* ^*11*^ *to quantify immunostaining. In order to compensate for moderate differences in expression levels between individual worms (carrying extrachromosomal arrays), we calculate percentages of fluorescence intensities in somata and synapses by considering the sum of fluorescence intensities (from somata and synapses) as being 100%* (Figures 8I and *S10B*+*D*).

#### General comments

Introduction: paragraphs do not connect to each other or flow logically. They are abrupt and need to be structured better.

Ordering figure panels can be better-the reader has to jump across multiple figure panels to look at data explained in one result section.

Figure panels need to be arranged/aligned properly and labeling is not proper in several panels.

*We have modified the revised manuscript accordingly.*

